# Wireless, Multimodal Monitoring of Organ Health Using 3D-Printed, Barbed, Bioresorbable Stretchable Microneedle Sensor Arrays

**DOI:** 10.1101/2024.07.16.603523

**Authors:** Xiangling Li, Shibo Liu, Jingshan Mo, Cheng Yang, Matthew Morales, Wei Ouyang

## Abstract

Comprehensive and continuous assessment of organ physiology and biochemistry, beyond the capabilities of conventional monitoring tools, can enable timely interventions for intraoperative complications like organ ischemia and nerve injuries and postoperative conditions such as organ dysfunction and transplant rejection. Here, we report a wireless implant with a 3D-printed, barbed, bioresorbable stretchable microneedle sensor array that offers multimodal monitoring of organ metabolism, oxygenation, and electrophysiology as well as spatiotemporal mapping of biomarkers across different organ regions. The development of a deformation-coupled 3D-printing technique enables 3D-programmable manufacturing of microneedles with monolithic backward-facing barbs, offering conformal yet robust 3D probing of organs with dynamic mechanics. Electrochemical functionalization of microneedle tips serves as a universal approach for localized sensing of physiological and biochemical parameters, enabling concurrent monitoring of up to 36 parameters (glucose, uric acid, oxygen, *etc.*) and spatiotemporal mapping of up to 32 sites. An electrically programmable self-destruction mechanism based on crevice corrosion and bioresorption eliminates the need for traumatic retrieval of microneedles. Demonstrations in clinically relevant complications like kidney ischemia and gut disorders in animal models highlight the broad applications of this device in intra- and postoperative monitoring.

---

Comprehensive and continuous monitoring of organ functions throughout the surgical course, from intraoperative evaluation to postoperative recovery, is essential for optimal patient outcomes^1, 2^. Intraoperative complications, arising from surgical procedures, anesthesia, medications, and individual patient susceptibilities, could lead to severe or even irreversible organ dysfunction that cannot be timely informed by traditional vital sign based intraoperative monitoring [*e.g.,* electroencephalography (EEG), electrocardiography (ECG), blood pressure (BP), oxygen saturation (SpO2)]^3–9^. Examples include acute kidney and liver injuries due to ischemia and/or nephron-/hepatotoxic agents, and bladder dysfunction due to accidental nerve damages^10–12^. On the other hand, capabilities in monitoring organ conditions in the postoperative phase, including potential surgical complications (*e.g.,* anastomotic leak after gastric surgeries)^13^, restoration of organ functions (*e.g.,* bladder control after partial cystectomy)^14^, transplant rejection (*e.g.,* the kidney and liver)^15–18^, and stress responses to surgery (*e.g.,* metabolic disorders, gut dysmotility)^19, 20^, is crucial to the recovery of patients. However, conventional methods like blood tests often lack specificity and show delayed responses to localized organ conditions^21, 22^. Radiological tests, while valuable, could only be performed intermittently in specialized facilities^23, 24^. Consequently, these conditions often remain undetected until after the development of severe outward symptoms, leading to substantial increases in morbidity and mortality^25^. These challenges underscore a critical, unmet clinical need for comprehensive and continuous assessment of organ functions to facilitate timely interventions during both intra- and postoperative phases^26, 27^.

Microneedles, being able to access organ parenchyma minimally invasively, holds great potential as ideal probes for organs^28, 29^. When electrochemically functionalized, microneedle sensors provide sensitive and specific measurement of a broad range of biochemical markers (*e.g.,* electrolytes, metabolites, and drugs) ^30–34^ in tissue and may directly integrate with electronics toward miniature, self-contained systems^35^, as demonstrated by recent works on interstitial fluid monitoring using skin-mounted microneedle patches^36^ and bladder monitoring using a microneedle-integrated catheter^37^. Nevertheless, the high modulus (∼GPa) of conventional microneedle arrays, typically fabricated from polymers like SU-8^38^, PMMA^39^, polyimide^40^, and polystyrene^41^, greatly exceeds that of organs (<1 MPa)^42^. This mechanical mismatch hinders conformal interfacing with the very soft organs, which often exhibit contractility (*e.g.,* the heart)^43^, motility (*e.g.,* the gut)^44^, and/or expandability (*e.g.,* the bladder)^45^, thereby compromising signal fidelity and burdening organs. Recent progresses in soft electronics lead to the development of a new class of microneedle arrays that are both flexible and stretchable^46, 47^. These novel microneedle arrays combine rigid microneedles with soft elastomers, allowing for intimate interfacing with dynamic organs. Despite these advancements, several key limitations remain for monitoring organ functions: (1) Existing stretchable microneedle arrays rely on sophisticated microfabrication processes^48^. Besides being costly and labor intensive and relying on sophisticated equipment, as a planar fabrication technology, microfabrication only allows for maximum microneedle lengths of ∼1 mm, as determined by substrate thickness, and does not offer programmability of microneedle lengths across the array. These limitations restrict the ability of microfabricated microneedle arrays to reach deep organ parenchyma and target different regions of varying depths. (2) Existing microneedle sensor arrays are typically electrically connected as a whole without addressability of individual microneedles and selectivity along the depth of microneedles, thereby losing 3D resolution^36^. This configuration does not allow for spatially resolved mapping of organ states across different functional regions and depths via individually addressed, localized microneedle sensors. (3) Conventional microneedles often show poor tissue adhesion, compromising the quality and stability of recording from organs with dynamic mechanics^37^. While the development of microneedles with swellable tips^49^, backward-facing barbs^50^, and tilted angles^51^, in the context of non-stretchable microneedle arrays, enhance tissue adhesion, these designs paradoxically complicate the removal process, increasing the risk of injury during extraction from sensitive organs. Therefore, microneedle arrays that seamlessly conform to and tightly adhere to dynamic organs, allow for precise probing and functional mapping of various organ regions and depths in a 3D manner, and ensure trauma-free removal post-use, represents an unrealized ideal for monitoring organs in intra- and postoperative settings.

Here we report a 3D-printed, barbed, bioresorbable Stretchable Microneedle Array for oRgan Tracking (SMART) that collectively addresses these limitations and its integration with wireless electronics into a fully implantable device for multimodal mapping of organ physiology and biochemistry in clinically relevant complications. Specifically, this work features the following advancements: (1) 3D-programmable microneedle arrays with monolithic backward-facing barbs through the development of deformation-coupled 3D printing for precise targeting of organ anatomy and strong tissue adhesion, eliminating the need for sophisticated 4D printing^50^ or magnetorheological drawing lithography^52^; (2) 3D-resolved functional mapping of organs through stretchable serpentine interconnects for addressing individual microneedles and selective exposure of microneedle tips for localized sensing; (3) Electrically programmable self-destruction to eliminate the need for microneedle retrieval through crevice corrosion of gold coatings and bioresorption of poly(lactic-co-glycolic acid) (PLGA) microneedles molded from 3D-printed structures; (4) Integration with wireless communication (Bluetooth-low-energy, BLE), a high-performance electrochemical frontend, and an analog switch matrix into a fully implantable device that allows for multimodal monitoring of electrolytes (*e.g.,* Na^+^, K^+^, pH), metabolites (*e.g.,* glucose (glu), uric acid (UA), lactic acid (LA)), tissue oxygenation, and electrophysiology and multisite mapping of up to 32 sites. Demonstrations in rodent models for monitoring ischemia and metabolic disorders in the kidney and gut highlight extensive capabilities of the device for comprehensive and continuous assessment of organ functions in clinically relevant complications.

## Results

### Design concepts and system features

The system consists of a fully implantable wireless device for monitoring diverse physiology and biochemistry of organs in intra- and postoperative settings, with real-time data wirelessly streamed to remote monitors (**Figs. 1a-b**). The wireless implant integrates a wireless electronics module and a SMART. The SMART seamlessly conforms to the organ surface, with microneedle sensors probing the organ parenchyma to measure diverse physiological and biochemical parameters of interest in a spatiotemporally resolved manner (**Fig. 1c**). The SMART anchors firmly to tissue through backward-facing barbs monolithically manufactured on microneedles to accommodate the dynamic mechanics of organs. Upon completion of monitoring, the barbed microneedles undergo electrically programmable self-destruction, through crevice corrosion and bioresorption processes, to eliminate the need for trauma-prone extraction from organs (**Fig. 1c**). As shown in **Fig. 1d** and **Supplementary Fig. 1**, the wireless electronics module (footprint: 21 mm×25 mm) consists of a BLE system-on-a-chip (SoC, nRF52832), an electrochemical frontend (AD5941), a 32-channel switch matrix (ADG732), a voltage buffer array (ADA4505), and a small lithium-polymer battery (65 mAh). The 32-channel switch matrix allows for multimodal monitoring of up to 32 amperometric biomarkers (*e.g.,* Glu, LA, UA, O_2_), multisite mapping of a single biomarker across up to 32 sites, or a combination of these 2 modes. The voltage buffer array allows for monitoring of multiple potentiometric biomarkers, such as K^+^, Na^+^, pH, and biopotential. The device features low power consumption of ∼0.5 mA in idle mode and ∼7 mA in active measurement mode (**Supplementary Fig. 2**). **Fig. 1e** and **Fig. 1f** show the photograph and exploded-view illustration of a 6×6 SMART, respectively. The SMART comprises a silicone (Ecoflex) base layer, a barbed microneedle array made of PLGA, a gold coating layer, a Parylene encapsulation layer that selectively exposes microneedle tips, and functionalization layers for the reference electrode (RE), the counter electrode (CE), and the working electrodes (WEs) with various sensing capabilities. **Fig. 1g** and **Fig. 1h** show completed devices with a 6×6 SMART and a 3×3 SMART resting on the palm, respectively, highlighting the miniature form factor of the device. The 6×6 SMART, measuring 1.5 cm×1.5 cm in the sensing area, may fit onto the surface of organs of humans or large animals. The 3×3 SMART, measuring 4 mm×4 mm in the sensing area, may fit onto the surface of organs of rodents. **Fig. 1i** shows a microscale computed tomography (micro-CT) image of a fully implanted device with a 3×3 SMART interfaced to the kidney of a rat, of which the electronics affixes to the abdominal wall by tissue adhesive. These results collectively demonstrate the form factor and functionality of the device suitable as an implant for comprehensive monitoring of organ states in various animal models.

**Fig. 1.**
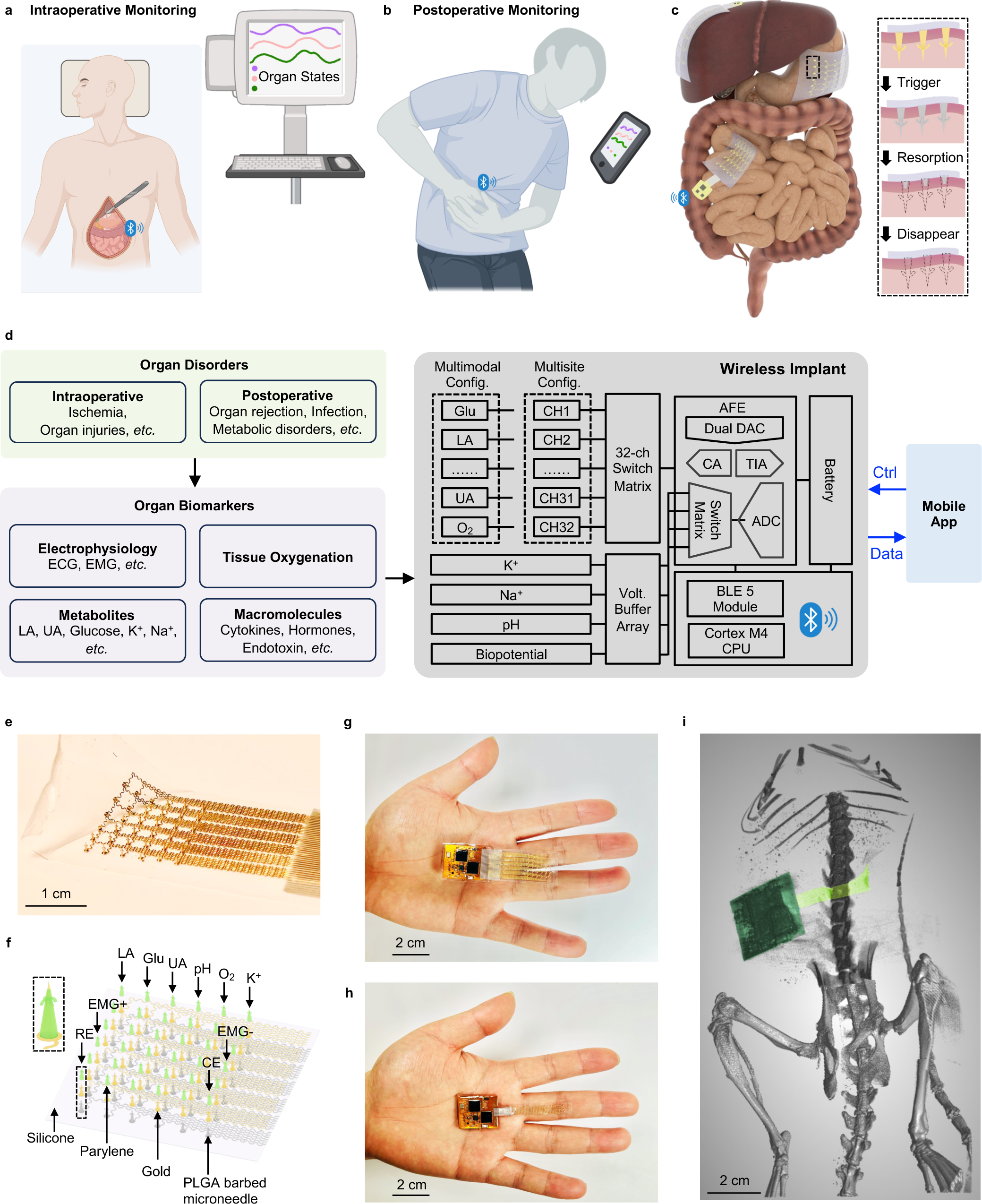
Design concepts and system features. **a,** Schematic illustration of the device for intraoperative monitoring of organ states. **b,** Schematic illustration of the device for postoperative monitoring of organ states. **c,** Schematic illustration of the SMART conformationally adhering to organs and electrically triggered bioresorption. **d,** Block diagram of the electronics and sensing functionalities of the wireless implant for multimodal and multisite monitoring of various organ disorders. **e,** Photograph of a 6×6 SMART. **f,** Exploded-view schematic illustration of the SMART and the different sensing modalities of the microneedles. **g,** Photograph of a device with a 6×6 SMART on the palm. **h,** Photograph of a device with a 3×3 SMART on the palm. **i,** Micro-CT image of a device with a 3×3 SMART implanted in an adult rat.

### Fabrication of the 3D-printed, barbed, bioresorbable SMART

**Fig. 2** and **Supplementary Fig. 3** illustrate the fabrication process. This process leverages 3D printing because it offers high programmability of microneedle diameters (≥100 μm) and lengths (from sub-millimeters to millimeters) across the array (**Supplementary Fig. 4**). However, as a layer-by-layer manufacturing technology, 3D printing does not support the direct manufacturing of microneedles with backward-facing barbs because printing a layer without the support of a preceding layer is impossible. To address this limitation, this work proposes a deformation-coupled 3D printing technique. The process starts with standard 3D printing of microneedles with horizontal barbs and serpentine interconnects (**Figs. 2a-b**). By default, the microneedles are 300 μm in diameter and 1000 μm in length, and the barbs are 80 µm in diameter and 400 µm in length. Next, a polydimethylsiloxane (PDMS) mold with cone-shaped patterns compresses the horizontal barbs into pre-designed backward-facing barbs (described in detail in the subsequent section), followed by curing of the 3D-printed resin under 405 nm ultraviolet (UV) light and removal of the PDMS mold (**Figs. 2c-d**). A double molding process using Ecoflex then transfers the barbed microneedle patterns from 3D printed resin to bioresorbable PLGA (**Figs. 2e-f**). Subsequently, sputtering of 10 nm Cr and 100 nm Au on the entire sample completes the metallization process (**Figs. 2g-h**). After activation of the contact surfaces by plasma treatment, a transfer printing process transfers and bonds the gold-coated PLGA structure to a thin film of silicone elastomers (Ecoflex, 500 μm thick) (**Figs. 2i-j**). Afterward, a PDMS slab screens the microneedle tips and self-stops upon reaching the barbs, followed by Parylene coating (3 μm) to encapsulate the microneedle bodies (**Figs. 2k-l**). A silicone sheet shields the substrate from Parylene coating. Upon removing the silicone covering sheet post-Parylene coating, spin coating of silicone on the substrate encapsulates the serpentine interconnects. Next, electrodeposition of gold nanoparticles (NPs, 20 nm in diameter) on the exposed microneedle tips enhances the electrical conductivity (**Figs. 2m-n**). Lastly, according to specific sensor functions, functionalization of individual microneedles with PEDOT:PSS, Prussian blue (PB), polyurethane (PU), ion-selective membranes (ISMs), enzymes, platinum NPs, polyvinyl butyral (PVB), and/or silver/silver chloride (Ag/AgCl) (**Supplementary Fig. 5**) completes the fabrication process. The completed SMART is highly stretchable, withstanding 20% stretching with maximum strain of 1.5% (**Fig. 2o**).

**Fig. 2.**
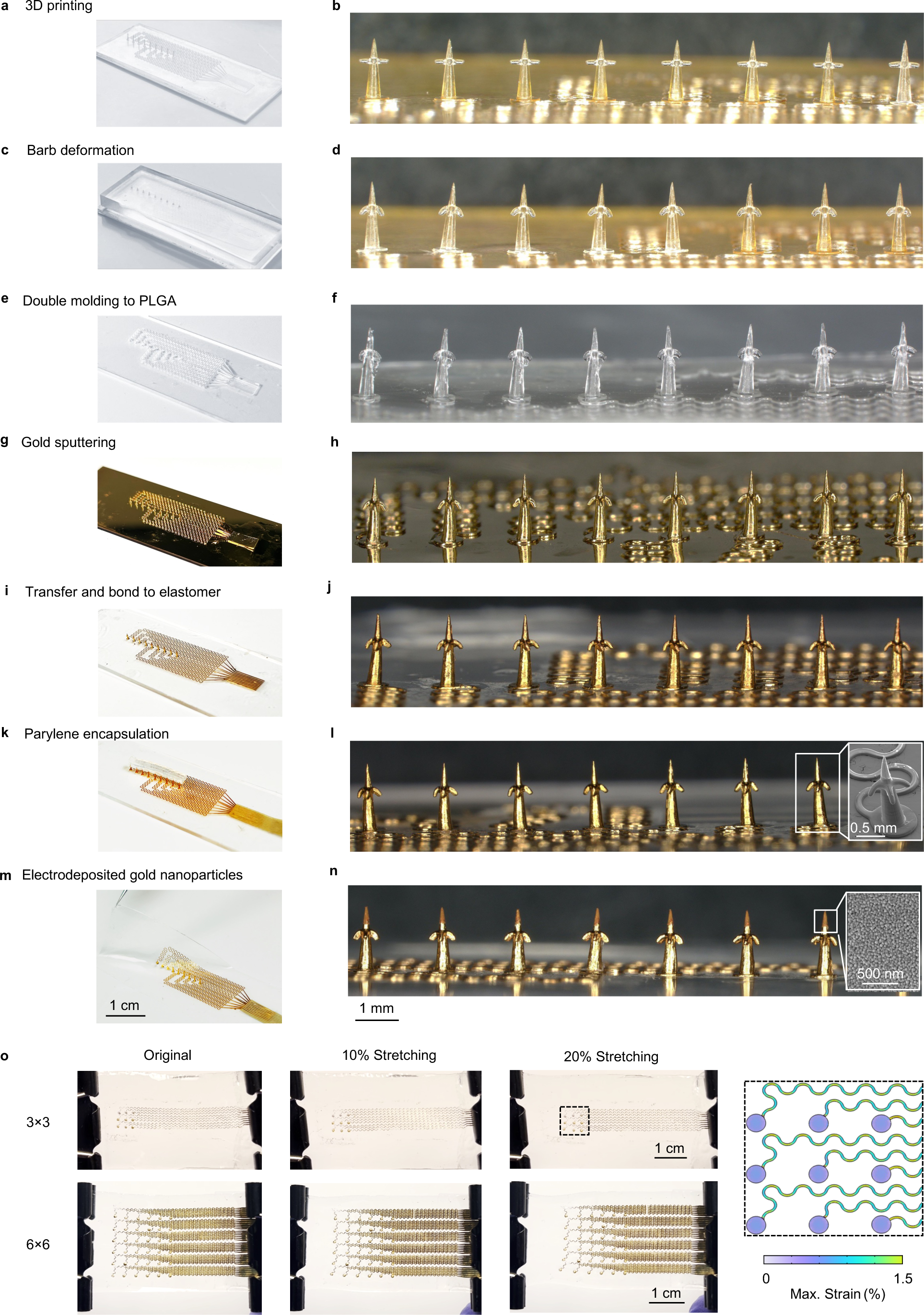
Photographs and micrographs of the SMART throughout the fabrication process. **a**, Photograph of the 3D-printed mold. **b**, Magnified view of the microneedles of the 3D-printed mold. **c**, Photograph of the barb deformation process using a PDMS mold. **d**, Magnified view of the microneedles after barb deformation. **e**, Photograph of the SMART after double molding to PLGA. **f**, Magnified view of the PLGA microneedles with barbs. **g**, Photograph of the SMART after gold sputtering. **h**, Magnified view of the PLGA microneedles with barbs after gold sputtering. **i**, Photograph of the SMART after transferring and bonding to elastomer. **j**, Magnified view of the PLGA microneedles with barbs after transfer. **k**, Photograph of the SMART with microneedles screened by a PDMS slab before Parylene coating. **l**, Magnified view of the PLGA microneedles with barbs after Parylene coating. **m**, Photograph of the completed SMART after electrodeposition of gold nanoparticles. **n**, Magnified view of the PLGA microneedles with barbs after electrodeposition of gold nanoparticles. **o**, Stretching performance of the SMART and numerical simulation.

### Tissue adhesion enhancement and electrically programmable bioresorption

Enhancing the tissue adhesion of microneedles critically relies on the monolithic fabrication of backward-facing barbs, as detailed in **Fig. 3a**. Following deforming horizontal barbs into backward-facing barbs using a PDMS mold, a molding process negatively transfers the patterns to Ecoflex. After temporarily bonding the Ecoflex mold to a glass slide to form an enclosed structure, infiltrating PLGA solution into the structure under vacuum and evaporating the solvent replicates the patterns from 3D-printed resin to PLGA. The low modulus of Ecoflex is crucial for the high-fidelity transfer of patterns with backward-facing barbs, as compared to the higher modulus of PDMS that distorts the transferred patterns (**Supplementary Fig. 6**). The PDMS mold for deformation consists of conical cavities with approximately the same heights as the microneedles and pre-designed angles to define the shape of the deformed barbs (**Supplementary Fig. 7a**). During compression, the barbs deform to the shape of the cones (**Supplementary Fig. 7b**). Cones with smaller angles results in more deformed barbs (**Supplementary Fig. 7c**). Simulation and experimental results indicate that the angle of the cone precisely defines the deformation of the barbs (**Supplementary Figs. 7d-i**). This method also allows for the monolithic manufacturing of multiple rows of barbs on a single microneedle (**Fig. 3b**). Furthermore, a single PDMS mold with individualized cone designs for microneedles of different dimensions allows for the manufacturing of barbs for the entire array in a single process with high yield (**Fig. 3c and Supplementary Fig. 8**). Tests in chicken breast, using barbs deformed by a 25° mold, indicate that microneedles with backward-facing barbs withstand strong pulling forces (**Fig. 3d**), which aligns well with simulation results (**Fig. 3e and Supplementary Fig. 9**). The critical pulling force, determined by applying different gravitational forces through weights (**Fig. 3f and Supplementary Fig. 9**), increases with the number of rows of barbs. For a 3×3 microneedle array, the critical pulling force increases from unmeasurable, to ∼0.02 N, ∼0.04 N, and ∼0.06 N for 0, 1, 2, and 3 rows of barbs per microneedle (**Fig. 3g**), respectively, which also agrees well with simulation results (**Fig. 3h**).

**Fig. 3.**
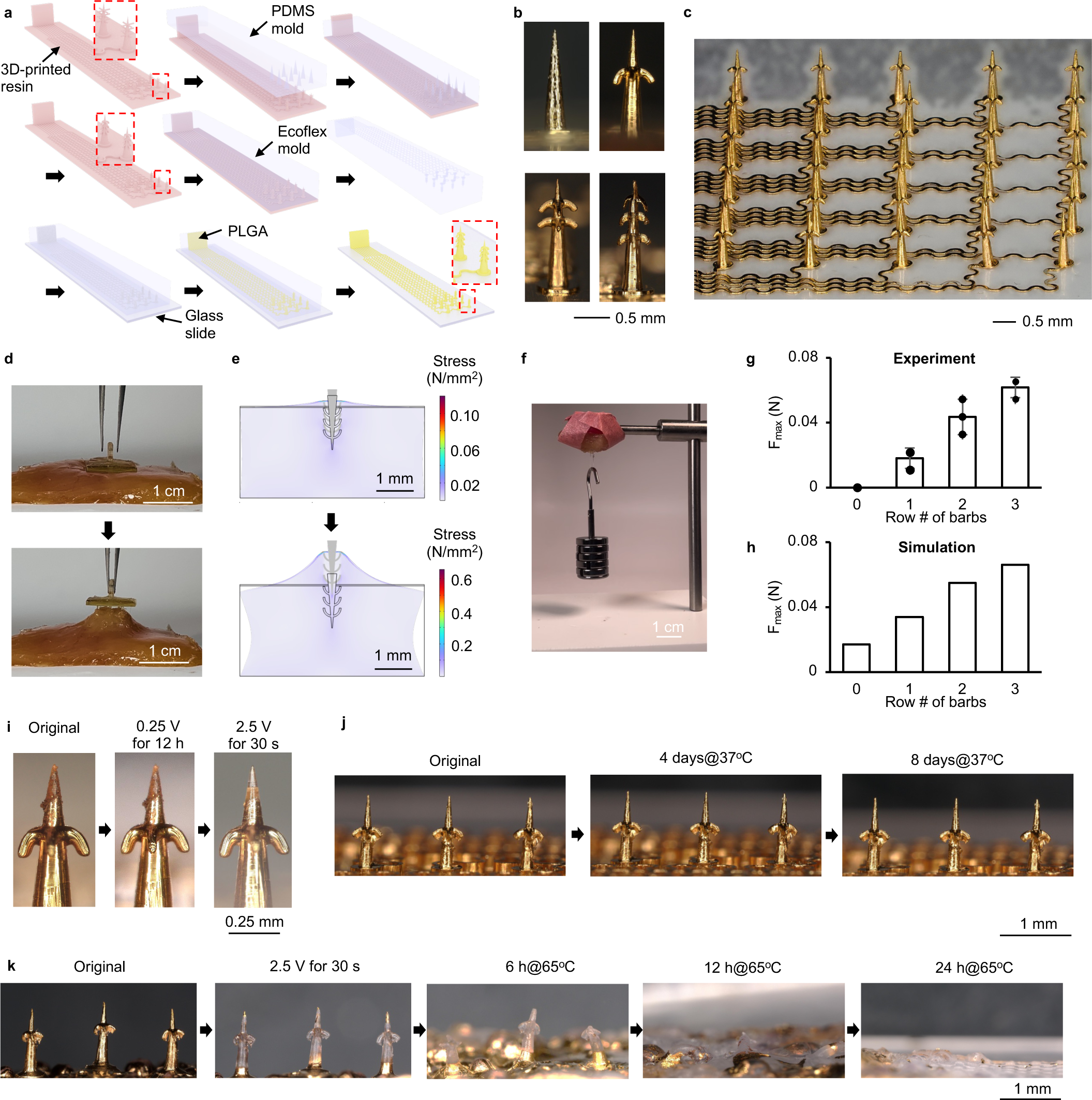
Tissue adhesion enhancement and electrically programmable bioresorption. **a,** Detailed schematic illustration of fabrication process of the barbed and bioresorbable SMART. **b,** Micrographs of barbed microneedles with different numbers of rows of barbs. **c,** Magnified view of a 6×6 microneedle array with barbs. **d,** Experimental images showing enhanced adhesion of barbed microneedles to a chicken breast sample. **e,** Numerical simulation results showing enhanced adhesion of barbed microneedles to tissue. **f,** An experimental image showing that barbed microneedles resist the pull force of five 10-g weights. **g,** Experimental results of the maximum pull force that barbed microneedles could withstand, which increases with the number of rows of barbs. **h,** Numerical simulation results of the maximum pull force that barbed microneedles could withstand, which agrees well with experiments. **i,** Micrographs showing that the functionalization layers do not degrade under long-term electrochemical measurements but disintegrate within 30 s under 2.5 V due to crevice corrosion. **j,** Time-lapse micrographs of microneedles in PBS at 37°C, indicating integrity of the microneedles. **k,** Electrically triggered disintegration of gold coatings followed by an accelerated bioresorption test in PBS at 65°C.

The SMART features an electrically programmable self-destruction mechanism to eliminate the need for trauma-prone retrieval of barbed microneedles from organs upon completion of monitoring. The microneedle sensors work stably during electrochemical measurements, showing no sign of degradation in the gold coatings after 12 h of amperometric measurement at 0.25 V (**Fig. 3i**). However, an overpotential exceeding 1.95 V triggers oxidative corrosion of the gold^53^. Subsequent mechanical crevices and exfoliations accelerate the disintegration of gold coatings. Consistent with literature^53^, the gold coatings at the exposed microneedle tips of the SMART completely disintegrate within 30 s at 2.5 V (**Fig. 3i**). Before the disintegration of the gold coatings, the PLGA microneedles and the gold coatings remain intact for over 10 days in phosphate buffered saline (PBS) at 37°C that simulates *in vivo* environment, indicating the robustness of the microneedle sensors (**Fig. 3j**). However, once electrically triggered, the PLGA microneedles start to degrade in organs through an FDA-approved, biologically safe hydrolysis process^54^. Using unencapsulated microneedles, accelerated degradation tests in PBS at 65°C suggest that the PGLA microneedles completely degrade within 24 hours (**Fig. 3k**), which corresponds to 5 days at body temperature (37°C) according to the Arrhenius equation^55^. These benchtop results demonstrate the feasibility of electrically programming the destruction of the microneedle sensors, thus eliminating the need for traumatic retrieval from organs in a secondary surgery.

### Microneedle functionalization for electrochemical sensing

The reported fabrication process offers high-yield and highly reproducible manufacturing of microneedle sensors, as demonstrated by consistent amperometric responses across a 6×6 array in glucose sensing (**Supplementary Fig. 10a**). The exposed tips of all the microneedles include a gold layer electrodeposited with Au NPs. Further microneedle-specific coatings yield different sensors. Electrodeposition of Pt NPs on a microneedle forms the counter electrode, and dip coating a microneedle in Ag/AgCl paste creates the reference electrode. For potentiometric sensing of ions (K^+^, Na^+^, and H^+^), sequential coating of a PEDOT:PSS ion-to-electron transducing layer and an ISM layer on AuNP-coated microneedles produces the working electrodes (**Fig. 4a**). The ISMs selectively allow the penetration of target ions, thereby creating an ion concentration gradient across the membrane and generating a trans-membrane potential indicative of the ion concentration. SEM shows the morphology of a K^+^-selective working electrode (**Fig. 4b**), indicating high uniformity of coating. Moreover, coating PVB on the Ag/AgCl reference electrode ensures a stable reference potential in solutions with different ionic strengths. The measured trans-membrane potential increases linearly with the K^+^ concentration from 1 mM to 32 mM (**Fig. 4c**), with a sensitivity of 69.8±2.63 mV/(lg[K^+^]), good linearity of R^2^= 0.996, and high reproducibility across three samples (**Supplementary Fig. 10b**). Similarly, in the case of Na^+^ sensing (**Fig. 4d**), as the concentration of Na^+^ increases from 5 mM to 160 mM, the measured potential increases linearly, with a sensitivity of 61.04±0.16 mV/(lg[Na^+^]), good linearity of R^2^= 0.999, and high reproducibility across three samples (**Supplementary Fig. 10c**). For pH sensing (**Fig. 4e**), as the pH value increases from 4 to 8, the measured potential increases linearly, with a sensitivity of −26.8±1.2 mV/(-lg[H^+^]), good linearity of R^2^= 0.994, and high reproducibility across three samples (**Supplementary Fig. 10d**). Furthermore, the continuous addition of interfering substances, such as NaCl, KCl, CaCl_2_, MgCl_2_, and NH_4_Cl, minimally affects the trans-membrane potential (**Supplementary Figs. 10e-f**), in contrast to the sensitive responses to target ions. These results collectively indicate the good sensitivity and specificity of the microneedle sensors for ion detection.

**Fig. 4.**
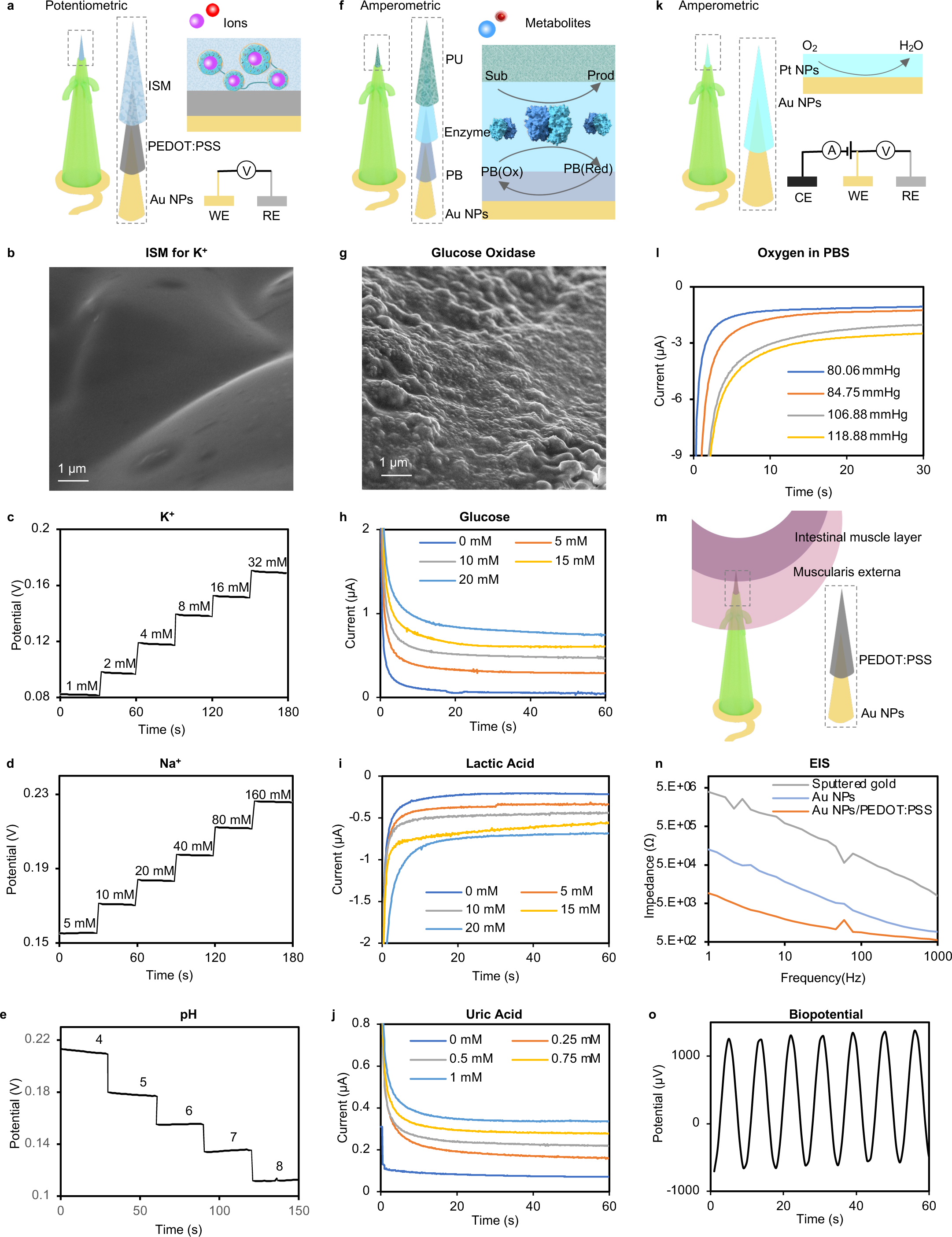
Sensing functionalities of the SMART. **a**, Schematic illustration of potentiometric sensors. **b**, SEM image of the K^+^-selective membrane coated on a microneedle tip. **c**, Potentiometric response of the K^+^ microneedle sensor to different concentrations of K^+^. **d**, Potentiometric response of the Na^+^ microneedle sensor to different concentrations of Na^+^. **e**, Potentiometric response of the pH microneedle sensor to different pH values. **f**, Schematic illustration of amperometric sensors. **g**, SEM image of glucose oxidase coated on a microneedle tip. **h**, Amperometric response of the glucose microneedle sensor to different concentrations of glucose. **i**, Amperometric response of the LA microneedle sensor to different concentrations of LA. **j**, Amperometric response of the UA microneedle sensor to different concentrations of UA. **k**, Schematic illustration of the oxygen sensor. **l**, Amperometric response of the oxygen sensor to different concentrations of dissolved oxygen. **m**, Schematic illustration of the EMG electrode. **n**, EIS characteristics of the EMG electrode. **o**, Sine waves recorded by the EMG electrode.

The monitoring of metabolic biomarkers, including Glu, LA, and UA, leverages enzyme-based amperometric sensors in a three-electrode electrochemical system (**Fig. 4f**). The electrons from enzyme-catalyzed oxidation of metabolites, efficiently transferred to the working electrode by electron mediators, generate an electrical current proportional to metabolite concentration. However, the low amplitude of the current resulting from the small sensing area of the microneedle tips limits the detection sensitivity and linear range. To address this limitation, PU serves as a diffusion-limiting layer at the outmost of the electrode to broaden the linear range of detection and isolate interfering chemicals^56^. At the same time, PB serves as an electron mediator to lower the overpotential applied to the electrode, thereby minimizing interference from other electroactive species, such as UA or ascorbic acid^57^. **Fig. 4g** shows the morphology of a microneedle electrode coated with glucose oxidase. Under a bias potential of 0.25 V, the microneedle glucose sensor detects glucose in the physiologically relevant range of 0 mM to 20 mM (**Fig. 4h**), with a sensitivity of 0.087±0.012 µA/mM, good linearity of R^2^= 0.9917, and high reproducibility across three samples (**Supplementary Fig. 10g**). In contrast to glucose-induced signals, sequential addition of 0.5 mM UA, 0.5 mM LA, 10 mM KCl, 0.01 mM vitamin C (VC), 1 mM NaCl, 0.5 mM UA, 0.5 mM LA, 10 mM KCl, 10 mM NaCl, 0.5 mM CaCl_2_, generates negligible currents (**Supplementary Fig. 10h**). Similarly, LA oxidase-coated and UA oxidase-coated microneedles serve as the sensors for LA and UA, respectively. Under a bias potential of −0.35 V, the current increases linearly with LA concentration in the range of 0 mM to 20 mM (**Fig. 4i**), with a sensitivity of 0.025±0.001 µA/mM, good linearity of R^2^= 0.9995, and high reproducibility across three samples (**Supplementary Fig. 10i**). Under a bias potential of 0.25 V, the current increases linearly with UA concentration in the range of 0 mM to 20 mM (**Fig. 4j**), with a sensitivity of 0.13±0.005 µA/mM, good linearity of R^2^= 0.9948, and high reproducibility across three samples (**Supplementary Fig. 10j**). Similar interference tests show that interfering chemicals generate negligible signals on the LA and UA sensors (**Supplementary Figs. 10k-l**), suggesting good selectivity of sensing. The measurement of tissue oxygenation, also known as oxygen partial pressure (PO_2_), utilizes the gold-standard Clark method^58^. Electrodeposited Pt NPs on AuNP-coated microneedles catalyzes the reduction of oxygen to generate an electrical current proportional to dissolved oxygen concentration (**Fig. 4k**). Under a bias voltage of −0.65 V, the amplitude of the current signal increases linearly with the concentration of oxygen in the PBS (**Fig. 4l** and **Supplementary Fig. 10m**), with a sensitivity of −0.037±0.006 µA/mmHg and good linearity of R^2^= 0.9995.

High-fidelity recording of biopotential critically relies on low impedance electrodes^59^. This work leverages a tri-layer coating of Au/AuNPs/PEDOT:PSS on microneedles to faithfully record organ electrophysiology (**Fig. 4m**). Electrochemical impedance spectroscopy (EIS) reveals electrical impedance of 7.8×10^5^ Ω, 2.7×10^3^ Ω, and 875 Ω at 100 Hz for microneedles with Au, Au/AuNPs, and Au/AuNPs/PEDOT:PSS coatings, respectively, highlighting nearly 3 orders of magnitude improvement by the tri-layer design (**Fig. 4n**). Benchtop testing suggests that the microneedle electrodes is capable of recording mV-scale sinusoidal signals in PBS (**Fig. 4l**), thereby meeting the needs of electromyography (EMG) recording (typical range: 0 – 10 mV) ^60^ in this work.

### Multimodal monitoring of organ health in rodent models

Demonstrations of multimodal monitoring of organ health involve implanting wireless implants with electrochemically functionalized 3×3 SMARTs in rodent models (adult Sprague-Dawley rats). Following the opening of the abdominal cavity via ventral midline or flank laparotomy, the SMART conformationally adheres to the surface of the target organ and establishes strong anchoring through the backward-facing barbs. The wireless electronics module affixes to the abdominal wall via tissue adhesive (**Supplementary Fig. 11**).

The first study focuses on organ ischemia, a common intraoperative complication that may result from low blood pressure, blood vessel injuries, and/or unsuccessful blood vessel connection in organ transplantation surgery^61^. This condition may also emerge post-surgery, such as in cases of organ transplant rejection^62^. The experiment simulates kidney ischemia through periodic occlusion of the renal artery and the renal vein using hemostatic forceps (**Fig. 5a**). The SMART probes the renal cortex and monitors renal oxygenation (PO_2_), LA, and UA (**Fig. 5b**). As the kidney undergoes 15-min cycles of ischemia and recovery, the device detects prompt decreases in PO_2_ to ∼29 mmHg upon ischemia and resumptions of PO_2_ to a baseline of ∼40 mmHg upon restoration of blood supply (**Fig. 5c**). Concurrent monitoring of renal LA and UA during these cycles, however, reveals steady increases in LA from ∼0.04 mM to ∼7 mM and UA from ∼0.03 mM to ∼0.33 mM over the course of 80 min. Furthermore, similar experiments in two separate rats yield comparable results (**Supplementary Fig. 12**). These findings might result from irreversible organ injury due to ischemia or reperfusion organ injury upon restoration of blood supply due to oxidative stress and inflammation^63^. The differential responses of different biomarkers to ischemia highlights the need for concurrent monitoring of multiple biomarkers, including oxygenation and biochemical markers, to ensure comprehensive assessment of organ health. To demonstrate the capability of spatiotemporally resolved monitoring of organ states, the 3×3 SMART maps renal oxygenation over 3×3 sites in the renal cortex during ischemia-recovery cycles (**Fig. 5d**). The device reveals spatial heterogeneity of oxygenation across different areas of the renal cortex and their temporal evolution upon the induction of hypoxia and recovery from hypoxia, showcasing the capability in high spatiotemporal resolution monitoring of organs.

**Fig. 5.**
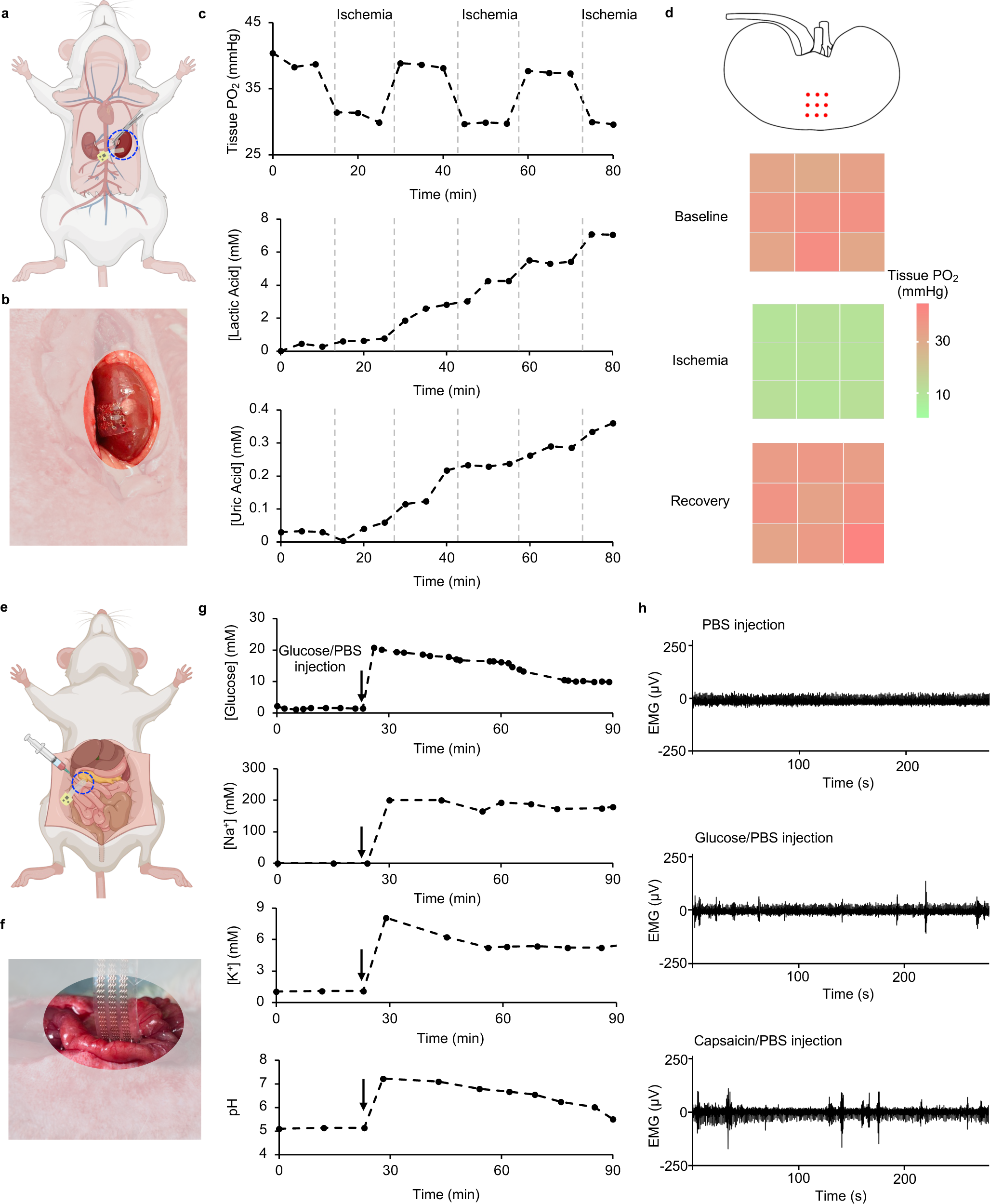
Organ monitoring in animal models. **a**, Schematic illustration of kidney monitoring using the device. **b**, Photograph of the SMART adhering to the kidney of a rat. **c**, Concurrent monitoring of kidney oxygenation, LA, and UA under periodical cycles of ischemia. **d**, Spatiotemporal mapping of kidney oxygenation under periodical cycles of ischemia. **e**, Schematic illustration of gut monitoring using the device. **f**, Photograph of the SMART encircling the small intestine of a rat. **g**, Concurrent monitoring of glucose, Na^+^, K^+^, and pH in the lumen of the intestine. **h**, EMGs of the intestine upon injection of PBS, glucose, and capsaicin.

The second study investigates postoperative gut disorders, such as nutrient absorption disorders, gut dysmotility, and anastomotic leaks, that may arise from gastrointestinal surgery, anesthesia and medications, and/or systemic inflammation and infection^64^. The SMART is capable of monitoring nutrient absorption within the lumen, tracking gut motility through EMG measurements on the intestinal wall, and detecting anastomotic leaks into the abdominal cavity, by adjusting the lengths of the microneedles. The SMART forms a cuff around a segment of the small intestine (jejunum), with the microneedles minimally invasively probing the lumen or the intestinal wall (**Figs. 5e-f**). Direct injection of electrolytes (PBS) and a nutrient (glucose) into the small intestine of anesthetized rats simulates food intake in this experiment. Upon injection, the device detects abrupt increases in the levels of glucose, Na^+^, K^+^, and pH that match those in the solution and their gradual decreases during absorption (**Fig. 5g**). Specifically, the glucose concentration in the lumen decreases from ∼20 mM to ∼9.9 mM within 1 hour.

Simultaneously, the Na^+^ concentration decreases from ∼200 mM to ∼180 mm and the K^+^ concentration decreases from ∼8.1 mM to ∼5.5 mM. The pH decreases from 7.1 to 5.5 during this process. Similar experiments conducted in a separate rat show comparable results (**Supplementary Fig. 13**). These results highlight the device’s capability in monitoring the gut’s absorption function. A separate set of experiments investigates (**Fig. 5h**) gut motility under different stimulations in anesthetized rats by using microneedles inserted into the intestinal wall without reaching the lumen. Following the initial injection of PBS, the gut does not exhibit notable myoelectric activities within a 5-min window. However, subsequent injection of PBS/glucose (20 mM) elicits significantly increased myoelectric activities within 5 min. In the second group receiving PBS/capsaicin (100 μM) injection after PBS injection, the gut also exhibits strong myoelectric activities within 5 min. These findings demonstrate the device’s potential in assessing gut dysmotility.

Histological (Hematoxylin and eosin, H&E) assessments of organ tissues (heart, liver, spleen, kidney, and intestine) in the aforementioned implantation groups indicate no identifiable inflammation responses such as immune cell aggregation associated with implantation after 2 weeks in comparison to the non-implanted control group (**Fig. 6**). These results suggest the biocompatibility of the device as an implant. Experimental studies of other implantable microneedle devices for organs (*e.g.* the brain) show minimal inflammatory responses and toxicity, as additional support for our claims of biocompatibility^53^.

**Fig. 6.**
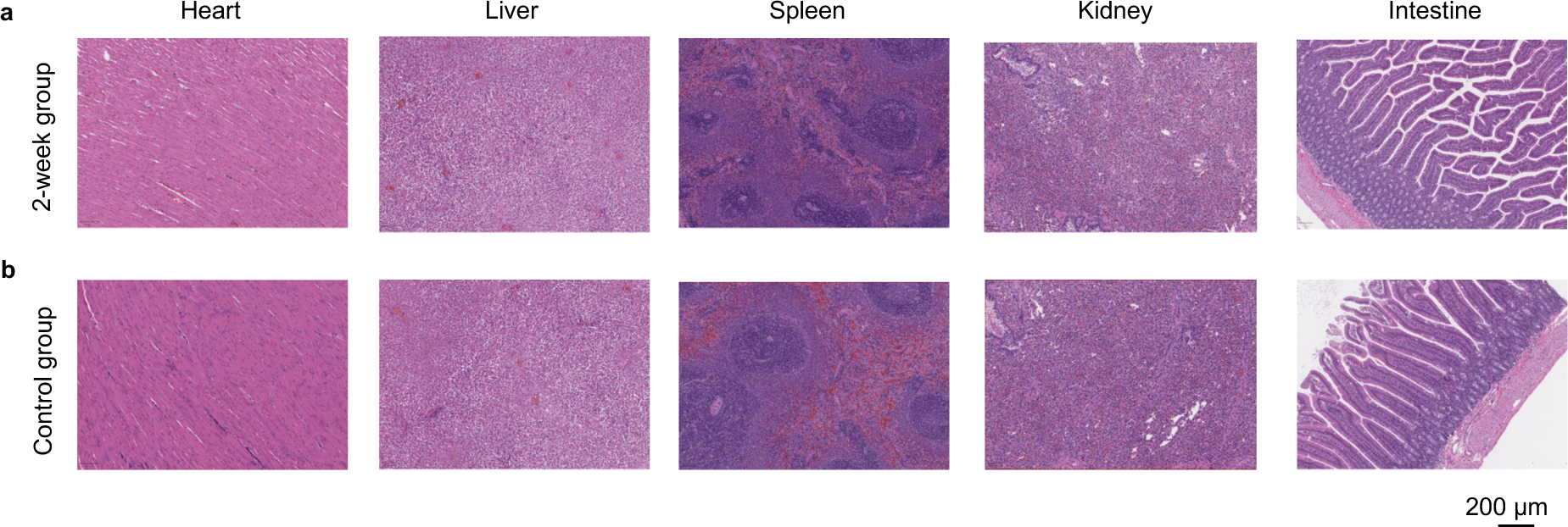
Biocompatibility study of the SMART. **a**, Representative images of H&E-stained tissue sections of the heart, liver, spleen, kidney, and intestine of rats 2 weeks after implantation of the SMART (2-week group). **b**, Representative images of H&E-stained tissue sections of the heart, liver, spleen, kidney, and intestine of healthy rats (control group).

## Discussion

This work presents a stretchable microneedle sensor array and its integration with wireless electronics for wireless, multimodal monitoring of organ health in intra- and postoperative settings. The development of a deformation-coupled 3D printing technique enables monolithic fabrication of microneedles with backward-facing barbs, thereby providing robust interface to organs. This method also offers high programmability of microneedle dimensions across the array, which, when combined with selective exposure of microneedle tips, enables spatiotemporally resolved mapping of organ states across different organ regions and depths. Double molding 3D-printed microneedles to PLGA provides a self-destruction mechanism through electrically triggered crevice corrosion of gold coatings and subsequent bioresorption of PLGA, thereby eliminating trauma-prone retrievable of barbed microneedles upon completion of monitoring. Electrochemical functionalization of the microneedles serves as a versatile approach for monitoring a broad range of biomarkers, including biochemicals (Glu, LA, UA, pH), tissue oxygenation, and electrophysiology. Demonstrations in monitoring clinically relevant complications in rodent models, including organ ischemia and gut disorders, underscore the crucial needs for continuous and comprehensive organ monitoring and the values of this device in this context. To completely avoid the need for a secondary surgery for device retrieval, the electronic components and battery for data measurement and wireless communication, in addition to the bioresorbable SMART, must be converted to bioresorbable formats using previously reported schemes in bioresorbable electronics^65, 66^. Future work integrating advanced electrochemical analysis techniques on microneedles would enable the real-time monitoring of other biomarkers, such as cytokines, drugs, and endotoxin, thereby opening up possibilities for investigating inflammation, infection, and drug metabolism in organs. Further chronic studies throughout the entire intra- and postoperative phases, in animal models of organ rejection, sepsis, and cancer, would offer invaluable insights into the pathophysiology and pharmacodynamics of important diseases.

## Methods

### Electrical components

The system was assembled on a double-layer fPCB [Cu (18 μm): Polyimide (PI, 75 μm): Cu (18 um)]. A low-dropout linear voltage regulator (ADP7112, Analog Devices Inc.) in a 6-pin wafer-level chip-scale package (WLCSP) (1.2 mm × 1 mm) converted the DC voltage to a constant voltage supply (3.3V) to power the system. A BLE SoC (nRF52832-CIAA, Nordic Semiconductor, Norway) in WLCSP packaging (3.0 mm × 3.2 mm) served as the CPU and Bluetooth communication module. The BLE SoC used a miniaturized (3.2 mm × 1.6 mm) ceramic 2.45 GHz antenna (2450AT18A100, Johanson Technology Inc.) for wireless communication. The potentiometric and amperometric circuitry consists of a voltage buffer array (ADA4505, Analog Devices Inc.), a 32-channel switch array (ADG732, Analog Devices Inc.), and an electrochemical analog front-end (AD5941, Analog Devices Inc.). The use of other passive components with 0201 (imperial) packaging minimized the overall size of the system.

### Fabrication of the SMART

The designed drawing was exported to a 3D printer (ELEGOO Saturn 3 UItra) and printed at a resolution of 10 μm per layer using an exposure time of 7 s. The microneedle array with barbed structures was obtained by using a PDMS mold with cone-shaped cavities to deform the horizontal barbs on the microneedles. Clamping them using binder clips and treating them under UV light at 405 nm for 3 min formed fully cured barb structures, followed by careful removal of the PDMS mold. Next, uncured Ecoflex (00-31) was cast to the 3D-printed structure. After vacuum removal of air bubbles and treatment at 70°C for 3 hours, the 3D-printed resin was carefully removed to obtain a negative mold of barbed microneedles and serpentines. The Ecoflex negative mold was then temporarily bonded to a glass slide to form an enclosed structure. It was then injected with polylactic acid (PLGA) solution (45 wt% in N-Methylpyrrolidone (NMP), Sigma-Aldrich) through the opening and vacuumed to remove bubbles. The mold was heated using a hot plate at 80°C for 24 hours with constant replenishment of PLGA solution to keep the mold filled. The mold was then transferred to a refrigerator at −20°C for 1 h. Next, quick removal of the mold yields a PLGA microneedle array with barbed structures and serpentines on a glass slide. Next, the microneedle array with barbed structures was metalized by sputtering 10 nm of Cr as the adhesive layer and 100 nm of Au as the conductive layer. A water-soluble tape (polyvinyl alcohol [PVA]) was patterned by using a CO_2_ laser to create circular holes matching the positions of the microneedles. This tape was applied downwards across the microneedle array onto the serpentines and the base of the microneedles and pressed for 1 minute to ensure strong adhesion. Gentle removal of the water-soluble tape transferred the microneedle array and serpentines from the glass slide to the water-soluble tape. The microneedle array on the water-soluble tape was flipped over to expose the back side, which was then plasma treated (Harrick Plasma, PDC001HP) for 5 minutes. The silicone elastomer thin film (Ecoflex 00-31; 500 μm thick) was obtained by spin-coating at 600 rpm. The silicone film was treated with a corona treater (BD-20AC, ETP Inc.) for 1 minute to form hydroxyl groups on the surface. Subsequently, the treated microneedle array was pressed onto the silicone elastomer thin film to form a tightly bonded structure. Finally, the water-soluble tape was washed off with water, followed by drying the microneedle array with an air gun.

### Device integration and encapsulation

An anisotropic conductive film (ACF) cable connected the fPCB and the SMART by hot pressing at 120°C for 15 seconds. A PDMS thin film (200 µm) was fabricated by spin-coating, followed by laser cutting (CO_2_) to form holes in the PDMS thin film that matched the positions of the microneedles. The PDMS thin film was applied downwards across the microneedle array onto the substrate and then gently pressed to squeeze out air bubbles. A PDMS slab was then inserted into the top of the microneedles until it reached the position of the barbs so that the depth of insertion into the PDMS could be precisely controlled. Subsequent deposition of a Parylene C layer (3 µm in thickness) encapsulates the entire device. The PDMS slab was then removed, leaving the tips of the microneedles exposed. The PDMS thin film was removed to eliminate the Parylene coating on the top side of the substrate. The Parylene coating on the bottom side of the substrate was also removed. A 500 µm thick layer of Ecoflex 00-31 was spin-coated on the substrate to complete the encapsulation of the serpentines. The fPCB was placed into a 3D-printed mold (30 mm×23 mm×3.6 mm). The mold was then completely filled with PDMS (10:1), followed by vacuum for 30 min to remove any air bubbles and curing at 75°C for 20 minutes. After removing the mold, the outer PDMS encapsulation layer of the fPCB was completed.

### Preparation of the Au/AuNP microneedle electrode

Au NPs were deposited on the tips of the microneedles after a 400-second plating reaction in a buffer solution containing 0.2 mM HAuCl_4_ (Sigma-Aldrich), 0.01 M HCl (Sigma-Aldrich), and 2 mM sodium citrate (Sigma-Aldrich) at a current of −1 mA.

### Preparation of the Au/AuNP/PtNP microneedle electrode

The Au/AuNP microneedle electrode was placed in a platinum-plating solution consisting of chloroplatinic acid (8 mM H_2_PtCl_6_ and 50 mM HCl) (Sigma-Aldrich). Pt NPs were deposited onto the Au/AuNP microneedle electrode after 200 seconds of electroplating reaction at a current of −1 mA.

### Preparation of the Au/AuNP/PB microneedle electrode

A Au/AuNP microneedle electrode was placed to a solution consisting of 0.25 mol/L potassium chloride (KCl), 0.05 mol/L anhydrous ferric chloride (FeCl_3_) and 0.05 mol/L potassium ferricyanide (K_3_[Fe(CN)_6_]) (Sigma-Aldrich). PB was deposited onto the Au/AuNP microneedle electrode after 120 seconds of electroplating reaction at a current of −0.5 mA to prepare the Au/AuNP/PB microneedle electrode.

### Preparation of the Au/AuNP/PEDOT:PSS microneedle electrode

The Au**/**AuNP/PEDOT:PSS microneedle electrode was obtained by immersing the Au/AuNP microneedle electrode in 30 ml of a mixture of 0.01 M (ethylene dioxythiophene monomer, EDOT) (Sigma-Aldrich) and 0.1 M sodium polystyrene sulfonate (NaPSS) (Sigma-Aldrich) and applying a constant current of 140 μA to the electrode for 200 seconds. The Au/AuNP/PEDOT:PSS microneedle electrodes were used for EMG recording.

### Preparation of the microneedle counter electrode and microneedle reference electrode

The Au/AuNP/PtNP microneedle electrode was used as the counter electrode. The fabrication of the Ag/AgCl reference electrode involved immersing gold-coated microneedle electrodes in Ag/AgCl paste (Sigma-Aldrich) to directly coat an Ag/AgCl layer on the electrode. Subsequently, the electrode was treated overnight in an oven at 80°C. For the preparation of the polyvinyl butyral (PVB) reference electrode, 79.1 mg of PVB and 50 mg of sodium chloride were dissolved in 1 ml of methanol. The electrode was then immersed in this solution for 3 hours. After removal from the solution, the electrode was treated at 90°C for 60 minutes to complete the fabrication of the PVB reference electrode.

### Preparation of glucose, uric acid, lactic acid, and O_2_ sensing microneedle electrode

(1) For glucose detection, a mixture consisting of glucose oxidase (50 mg/mL, Sigma-Aldrich), bovine serum albumin (BSA, 80 mg/ml, Sigma-Aldrich), and glutaraldehyde (2.5 wt% in phosphate-buffered saline (PBS), Sigma-Aldrich) was prepared in PBS at a volume ratio of 1:5:2. The Au/AuNP/PB microneedle electrodes were drop-coated three times in the glucose oxidase precursoe solution described above. Following overnight drying, the electrodes were immersed in a polyurethane (PU) solution (0.6 g of PU dissolved in 10 g of a 98:2 mass ratio tetrahydrofuran/ dimethylformamide solution, Sigma-Aldrich). Finally, air drying for 12 hours at ambient temperature of the resultant Au/AuNP/PtNP/GOx microneedle electrodes completed the fabrication process. (2) For uric acid detection, uric acid oxidase (15 mg/ml, Sigma-Aldrich), BSA (80 mg/ml, Sigma-Aldrich), and glutaraldehyde (2.5 wt% PBS, Sigma-Aldrich) were combined in PBS at a volume ratio of 1:5:2. The Au/AuNP/PB microneedle electrodes were processed using the dip-coating technique in the uric acid oxidase precursor solution, followed by overnight drying and immersion in PU solution (0.6 g of PU dissolved in 10 g of a 98:2 mass ratio tetrahydrofuran/ dimethylformamide solution, Sigma-Aldrich). Finally, air drying for 12 hours at ambient temperature of the resultant Au/AuNP/PB/Uricase microneedle electrodes completed the fabrication process. (3) For lactate sensing, lactate oxidase (30 mg/ml, Sigma-Aldrich), BSA (80 mg/ml, Sigma-Aldrich), and glutaraldehyde (2.5 wt% PBS, Sigma-Aldrich) were dissolved in PBS at a volume ratio of 1:5:2. The Au/AuNP/PB microneedle electrodes underwent the dip-coating process in the lactate oxidase solution for three times, followed by overnight drying and immersion in PU solution (0.6 g of PU dissolved in 10 g of a 98:2 mass ratio tetrahydrofuran/ dimethylformamide solution, Sigma-Aldrich). The electrodes, designated as Au/AuNP/PB/LOx microneedle, were subsequently dried for 8 hours at room temperature. (4) The Au/AuNP/PtNP microneedle electrode was used as the O_2_ electrode based on the Clark oxygen sensing method ^58^.

### Preparation of Na^+^, K^+^, and pH sensing microneedle electrodes

The preparation of the Na^+^-selective membrane cocktail was conducted using a composition that included Na ionophore X (1% w/w, Sigma-Aldrich), sodium tetrakis[3,5-bis(trifluoromethyl)phenyl] borate (Na-TFPB, 0.55% w/w, Sigma-Aldrich), polyvinyl chloride (PVC, 33% w/w, Sigma-Aldrich), and dioctyl sebacate (DOS, 65.45% w/w, Sigma-Aldrich). A quantity of 200 mg of this cocktail was solubilized in 1,320 μL of tetrahydrofuran (THF) under agitation for 30 minutes. For the K^+^-selective membrane, the formulation comprised valinomycin (2% w/w, Sigma-Aldrich), potassium tetrakis[phenyl]borate (NaTPB, 0.5% w/w, Sigma-Aldrich), PVC (32.7% w/w), and DOS (64.8% w/w). Here, 200 mg of the mixture was dissolved in 700 μL of cyclohexanone to fabricate the membrane solution. ISMs for Na^+^ and K^+^ were fabricated through a dip-coating process of the corresponding electrodes into their respective membrane cocktails, followed by air-drying at ambient temperature overnight. The pH-sensing coating was synthesized by dissolving 20 mg of polyaniline emeraldine salt in 20 mL of dimethyl sulfoxide (DMSO, Sigma-Aldrich). To achieve complete dissolution, the mixture was stirred continuously for 24 hours. The microneedle electrodes were then coated by dip-coating in the polyaniline emeraldine solution and dried for 2 hours at 60°C, thus forming a polyaniline emeraldine base membrane on the gold electrode surface. Subsequently, the membrane was exposed to a 1 mL solution of hydrochloric acid (HCl, 1 mol/L, Sigma-Aldrich) within a vacuum chamber for 6 hours to ensure thorough protonation.

### Surface morphology characterization of sensing electrodes

The surface morphology of the microneedle sensing electrodes was imaged by a benchtop scanning electron microscope (SEM, Helios 5 CX DualBeam, Thermo Fisher Scientific). Micrographs of the samples were taken using an optical microscope (VHX-7000N, Keyence).

### Electrochemical characterization of microneedle sensing electrodes

The microneedle electrodes (working electrodes) used for biochemical assays were characterized using an electrochemical workstation (PalmSens 4). The selectivity of the electrochemical sensors for glucose, lactate, uric acid, O_2_, sodium ions, potassium ions, and pH was evaluated separately by sequentially adding certain concentrations of interfering substances to the solution. For the ion sensors, the electrochemical workstation characterized the open-circuit potential of the two-electrode system, where a microneedle Ag/AgCl electrode was used as the reference electrode during the test. The open-circuit potential was recorded for different concentrations of the substance to be measured. A series of ion solutions with different concentrations were prepared, in which deionized water was used as the solvent. The test was paused for 30 seconds each time the concentration of the substance changed to allow for sufficient mass transfer in the solution. Interference studies were carried out by adding 10 mM NaCl, 10 mM KCl, 0.5 mM CaCl_2,_ or 5 mM NH_4_Cl dissolved in deionized water, respectively. For the metabolite sensors, a three-electrode system (working, counter, and reference electrodes) was used, where the amperometric sensors were applied bias potentials of 0.25V, −0.35 V, and 0.25 V for glucose, lactate, and uric acid, respectively. A microneedle Au/AuNP/PtNP electrode was used as the counter electrode, and a microneedle Ag/AgCl electrode was used as the reference electrode during the test. Interference studies were performed by adding 1 mM UA, 0.5 mM LA, 0.01 mM VC, 10 mM NaCl, 0.5 mM CaCl_2_, or 10 mM KCl dissolved in deionized water, respectively. The effect of the material on the overall sensor performance was assessed by measuring the electrochemical properties of electrodes with different material structures. The sensitivity, detection selectivity, and detection range of each electrode were analyzed separately. For the oxygen sensor, a three-electrode system (working electrode, counter electrode, and reference electrode) was used with a bias potential of −0.65V applied. A microneedle Au/AuNP/PtNP electrode was used as the counter electrode and a microneedle Ag/AgCl electrode was used as the reference electrode during the tests. Dissolved oxygen concentration in PBS was measured using the Digital Tester Smart Bluetooth Dissolved Oxygen Meter (RMR-6NW-2QU355, Buiishu). The concentration of oxygen in PBS was regulated by continuous passage of air or heating.

### Finite element analysis

The deformation process and tissue adhesion behavior of backward-facing barbs and the stretchability of the SMART were numerically investigated by the solid mechanics module of COMSOL Multiphysics. For the deformation process, a two-dimensional symmetric model was created as shown in Supplementary Fig. 7. A line segment was defined as the contact surface, with angles of 15°, 25°, and 35° relevant to the y-axis. The contact line underwent a fixed displacement to simulate the deformation process. The deformation and stress distribution of the barbs after contact were computed. To characterize tissue adhesion of the barbs, the microneedle was treated as a rigid body and the tissue was set as a linear elastic material. The main boundary of the model was set such that the bottom side was fixed, and a pair of equal tension forces was applied on both sides to simulate the tissue tension, and a gradually increasing force was applied on the upper part of the microneedle to simulate the extraction process. For modeling the stretchability, a simplified 3D model was used. A thin rectangle of 20 × 55 × 0.4 mm was defined as the flexible substrate and set as a hyperelestic material. The PLGA interconnects were characterized by a density of 1530 kg/m^3^, a Young’s modulus of 32.0 MPa, and a Poisson’s ratio of 0.12^67^.

### Animal Experiments

Experimental protocols were approved by the Institutional Animal Care and Use Committee of Dartmouth College. All animals were handled following the guidelines for the care and use of laboratory animals described by the Institutional Animal Care and Use Committee of Dartmouth College (Protocol # 00002330). Male Sprague-Dawley rats weighing 250-300 g (Animal Center of Dartmouth Hitchcock Medical Center) were used for experiments. Rats were housed in a climate-controlled room with a 12-hour light/12-hour dark cycle and fed *ad libitum*.

### Kidney monitoring using the SMART

Sprague-Dawley rats were anesthetized using an anesthesia system (Isotec 4, SurgiVet, United States) with 2% isoflurane gas delivered via a nose cone. The rats were placed on a thermostatic heat pad to maintain a constant body temperature throughout the procedure. Before the experiment, hair removal cream (VEET hair removal cream, Reckitt Benckiser) was used to remove hair from the abdomen of the rats. Subsequently, laparotomy was performed via a ventral midline or flank approach to open the abdominal cavity. The kidney was located, and a SMART was wrapped around the kidney and pierced into the renal cortex. Ischemia was induced every 15 minutes using hemostatic forceps to occlude the renal artery and the renal vein. Electrochemical signals were recorded every 5 min. Finally, a 3×3 SMART was used to map the spatiotemporal changes in renal oxygen levels using the same ischemia simulation method.

### Gut monitoring using the SMART

Sprague-Dawley rats were anesthetized using an anesthesia system (Isotec 4, SurgiVet, United States) with 2% isoflurane gas delivered via a nose cone. The rats were placed on a thermostatic heat pad to maintain a constant body temperature throughout the procedure. Before the experiment, hair removal cream (VEET hair removal cream, Reckitt Benckiser) was used to remove hair from the abdomen of the rats. Subsequently, laparotomy was performed via a ventral midline approach to open the abdominal cavity. A segment of the small intestine was extruded, and a SMART was wrapped around the jejunum. The microneedle electrodes minimally invasively pierced into the lumen of the intestine. Electrochemical signals for glucose, Na^+^, K^+,^ and pH were recorded at 5 min/10 min intervals, and at the 20th min of the experiment, 2 mL of 20 mM glucose/PBS solution was injected to study intestinal absorption. Subsequent recordings were made until the 90th minute. After the completion of the experiment, the animals were euthanized. For myoelectricity experiments in the intestine, the microneedles were located within the intestinal wall without puncturing the intestine. Two sets of experiments were conducted, in which 1 mL of PBS was injected first, followed by injection of 1 mL of 20 mM glucose/PBS solution. In the other set of experiment, 1 mL of PBS was injected first, followed by injection of 1 mL of capsaicin/PBS solution (100 uM).

### Microscale CT imaging

CT imaging was performed using a preclinical microPET/CT imaging system (X-RAY source) with “medium” magnification, 33 µm focal spot, 1 × 1 binning, 720 projection views, and 100 ms exposure time. The 3D image was reconstructed using the software Dragonfly.

### Reporting Summary

Further information on research design is available in the Nature Research Reporting Summary linked to this article.

## Data availability

The main data supporting the results in this study are available within the paper and its Supplementary Information. Source data for the figures will be provided with this paper.

## Acknowledgements

We acknowledge the startup fund to W.O. from Thayer School of Engineering, Dartmouth College.

## Author contributions

X.L. and W.O. conceived the ideas and designed the research. X.L. developed the sensors. X.L. and M.M. manufactured and tested the sensors. S.L. designed and manufactured the electronics. J.M. and C.Y. performed the finite element simulation. X.L. performed the sensor characterizations and animal experiments. X.L. and W.O. wrote the manuscript. All authors reviewed and commented on the manuscript.

## Supplementary Information

**Supplementary Fig. 1.**
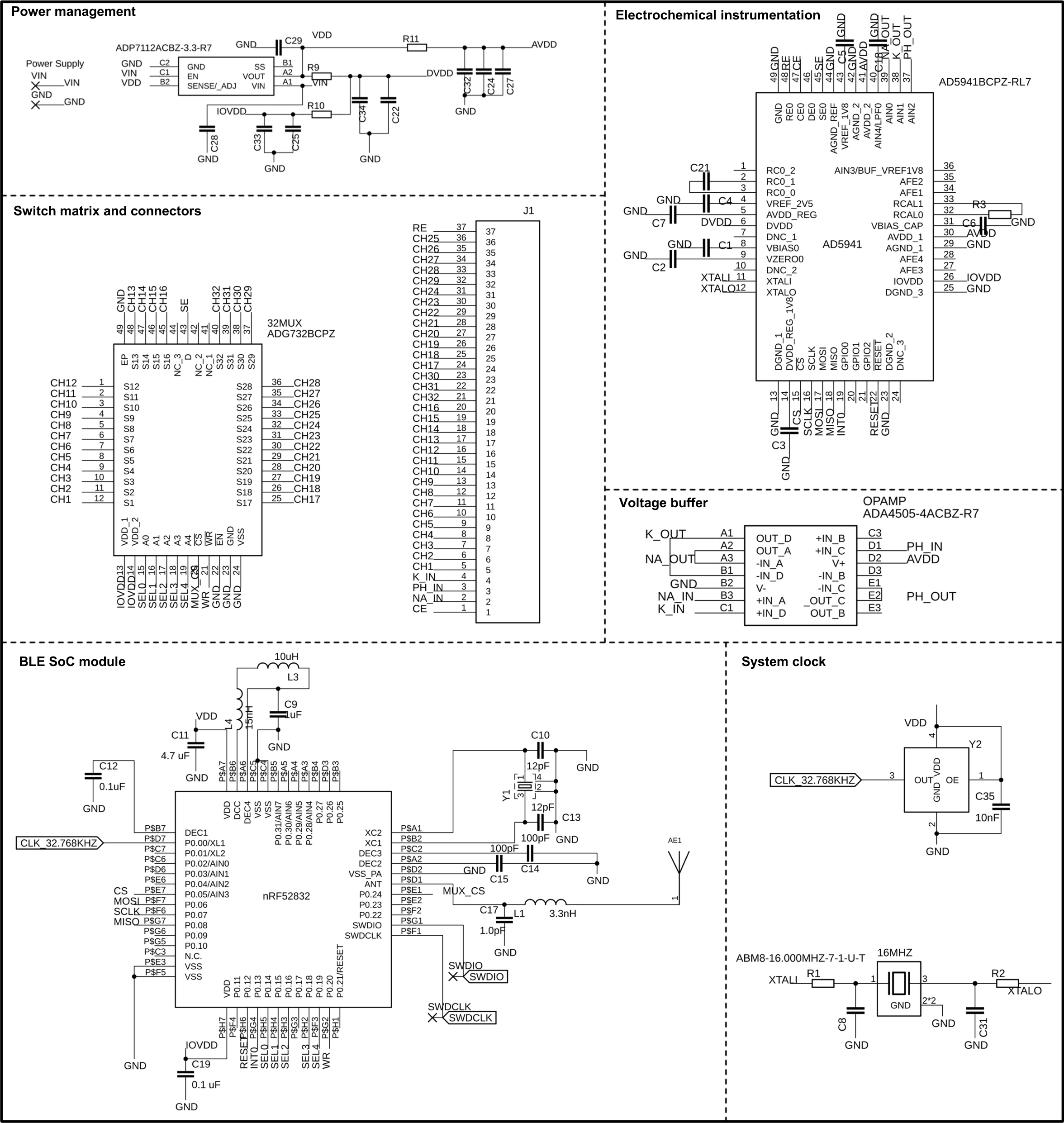
Electrical schematic of the wireless implant.

**Supplementary Fig. 2.**
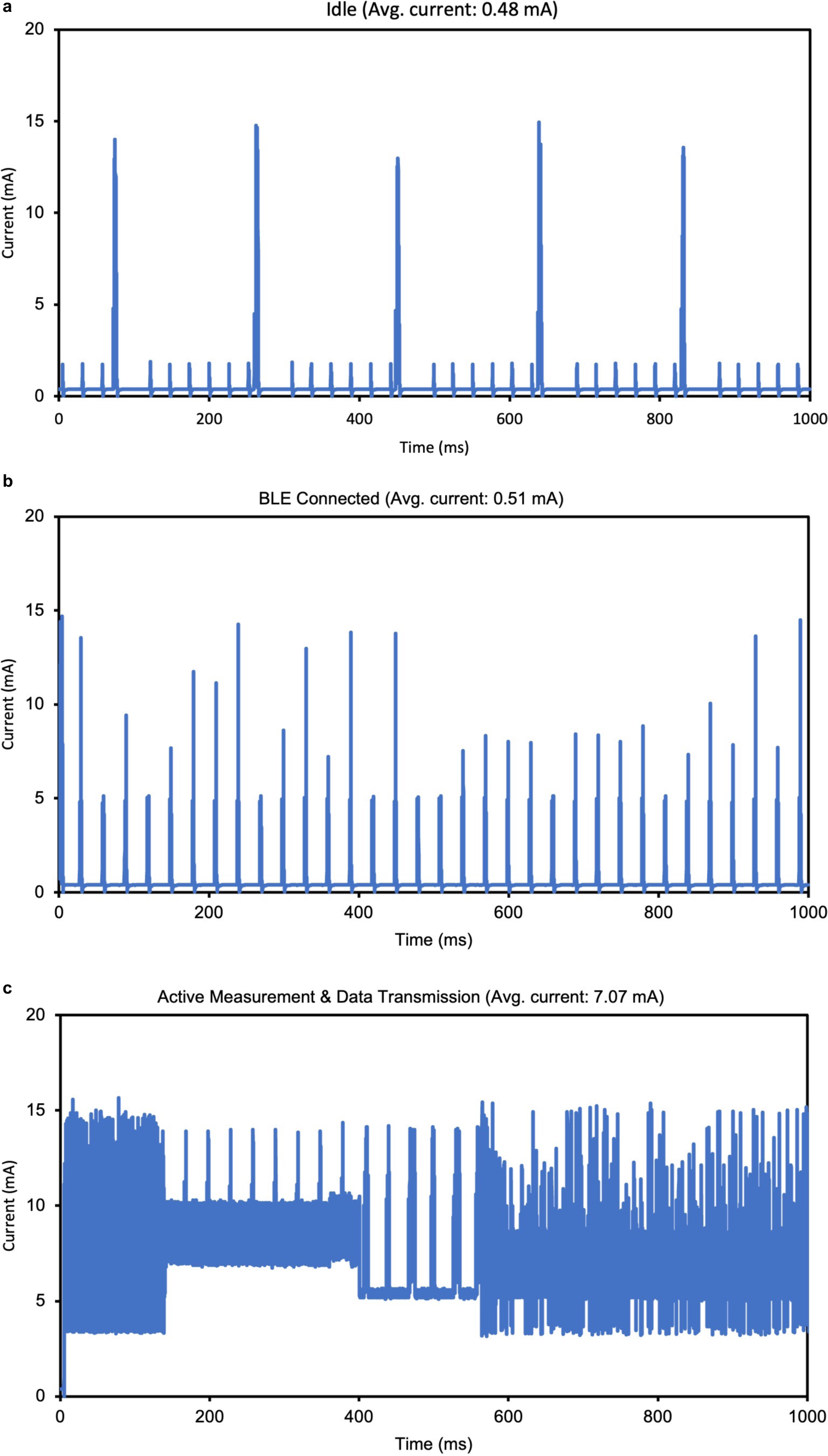
Current consumption of the wireless implant under different modes. **a**, idle mode. **b**, BLE connected mode. **c**, Active measurement and data transmission mode.

**Supplementary Fig. 3.**
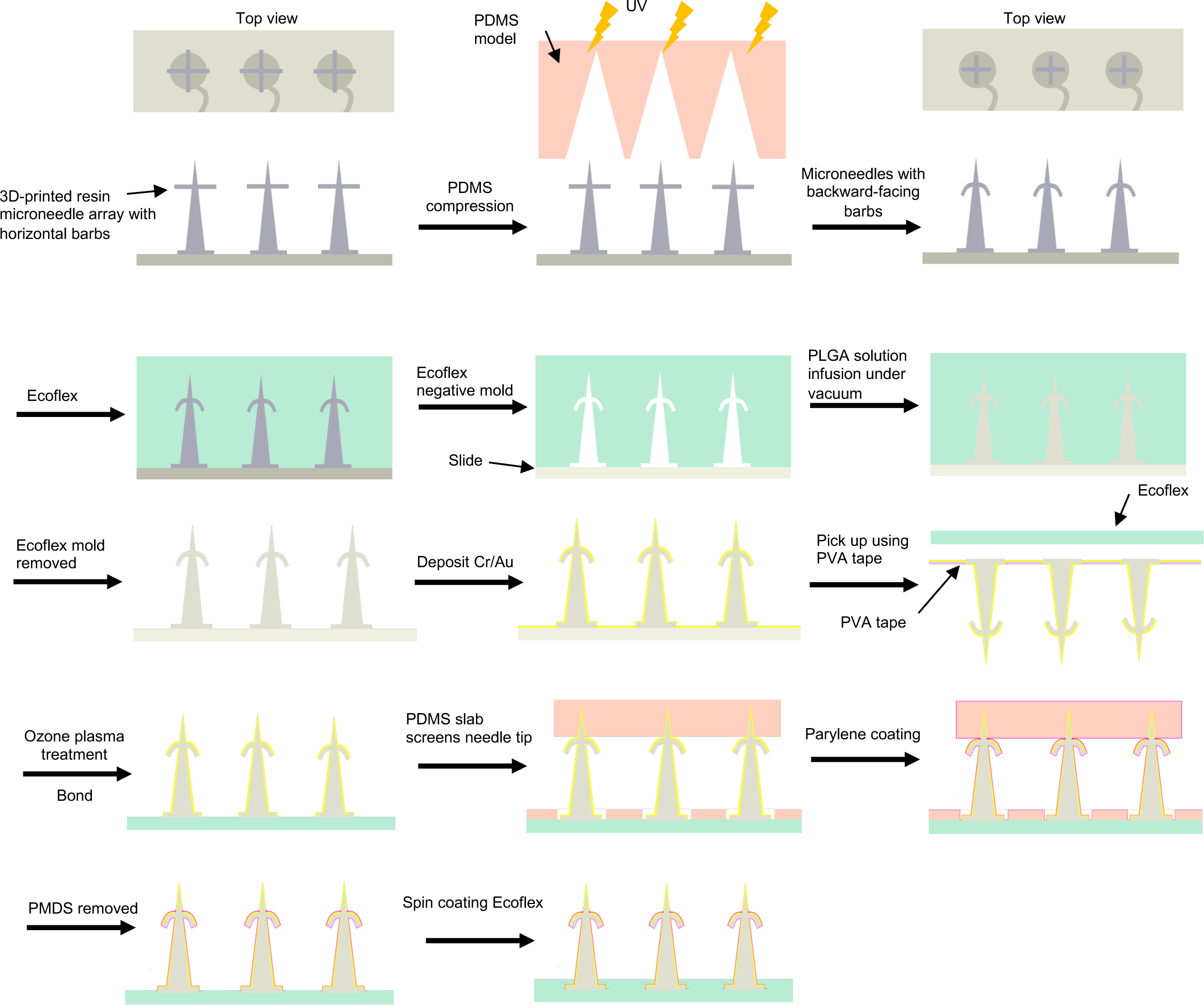
Schematic illustration of the fabrication process.

**Supplementary Fig. 4.**
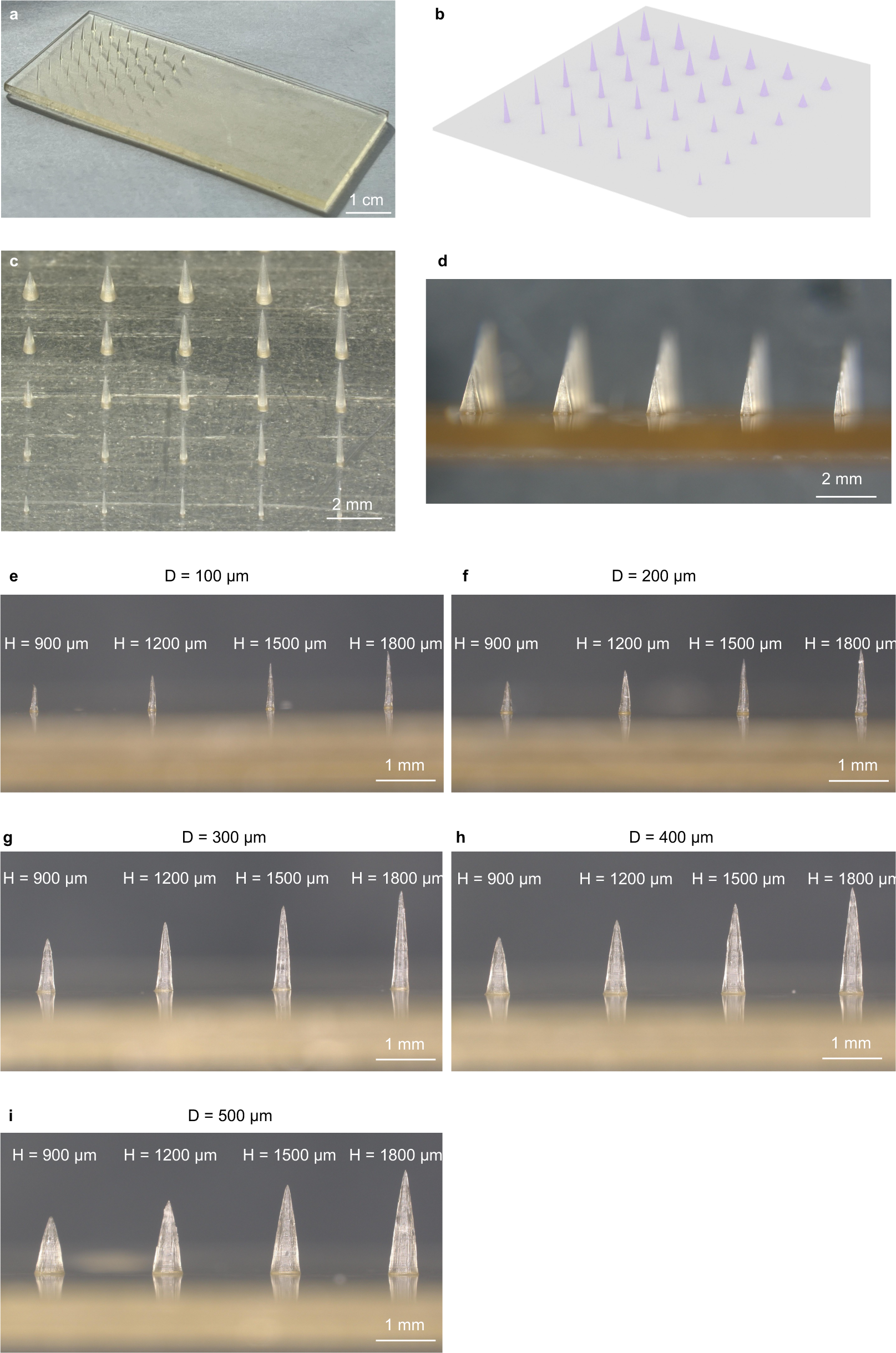
Highly programmable manufacturing of microneedles of different dimensions by 3D printing. **a,** A 3D-printed microneedle array with microneedles of diverse diameters and lengths. **b,** 3D rendering of the microneedle array. **c,** Top-view of the microneedle array. **d,** Side view of the microneedle array. **e-i,** Microneedles of different lengths and diameters of **e,** 100 μm. **f,** 200 μm. **g,** 300 μm. **h,** 400 μm. **i,** 500 μm.

**Supplementary Fig. 5.**
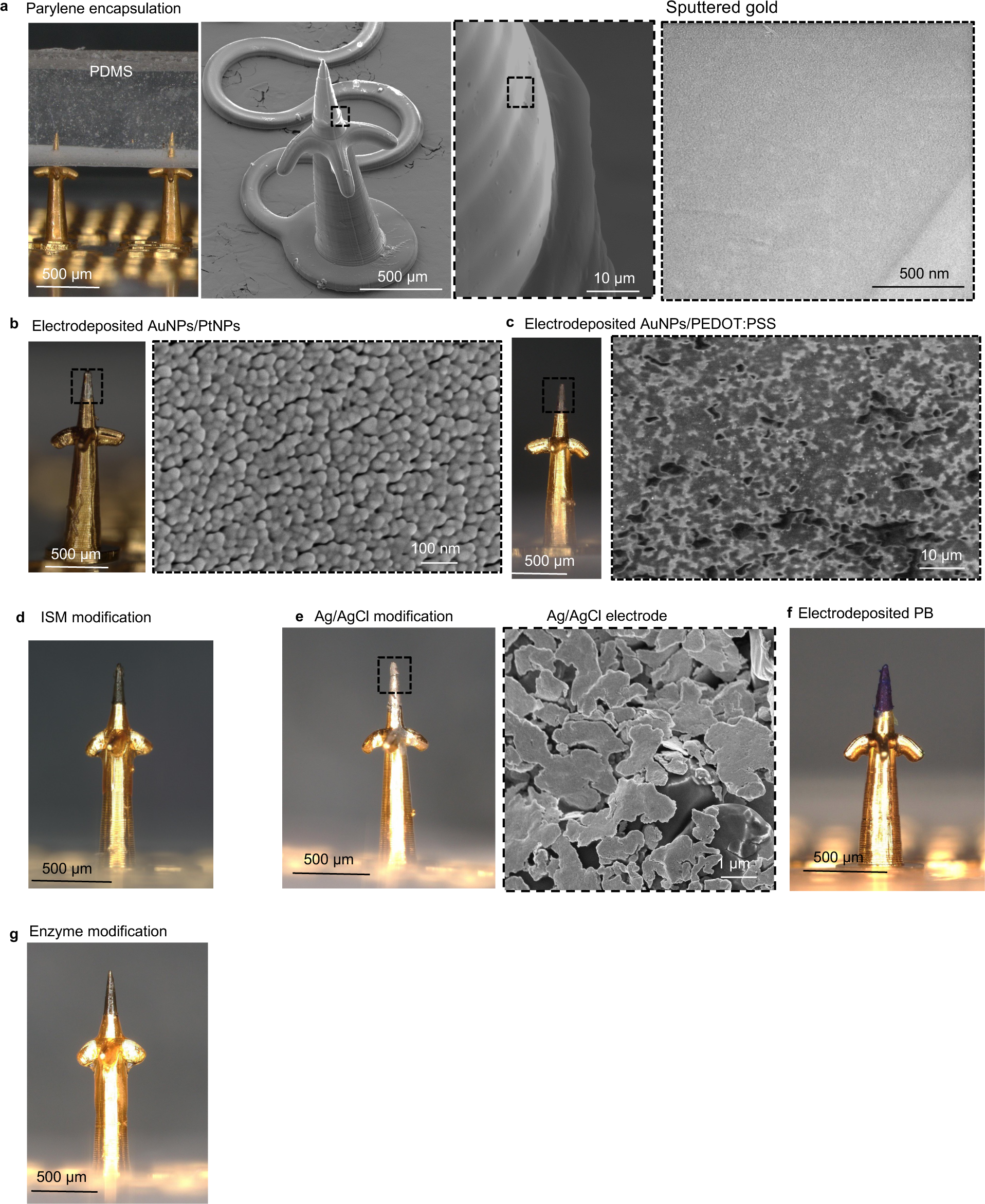
Micrographs of microneedles after coating different functionalization layers. **a,** After Parylene encapsulation. **b,** After electrodeposition of AuNPs/PtNPs. **c,** After electrodeposition of AuNPs/PEDOT:PSS. **d,** After coating of an ISM. **e,** After coating of Ag/AgCl. **f,** After coating of PB. **g,** After coating of an enzyme.

**Supplementary Fig. 6.**
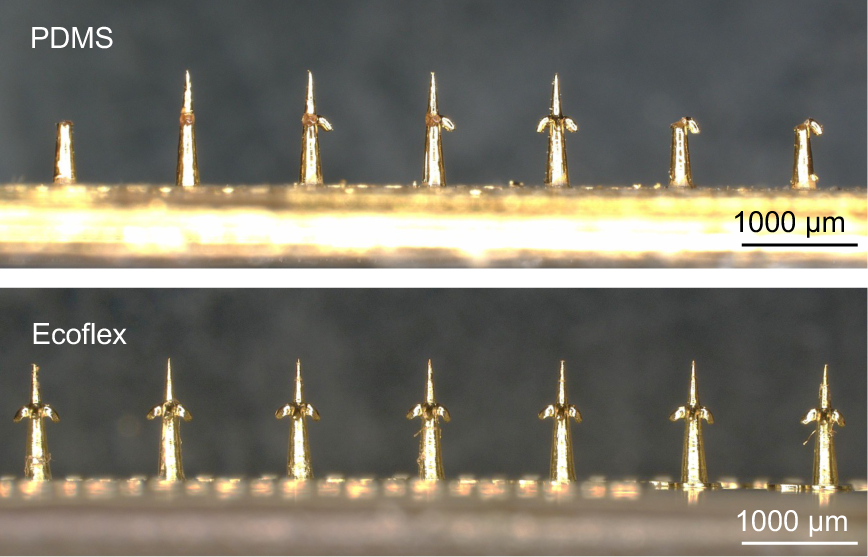
Comparison of the microneedle array after double molding using PDMS and Ecoflex, respectively.

**Supplementary Fig. 7.**
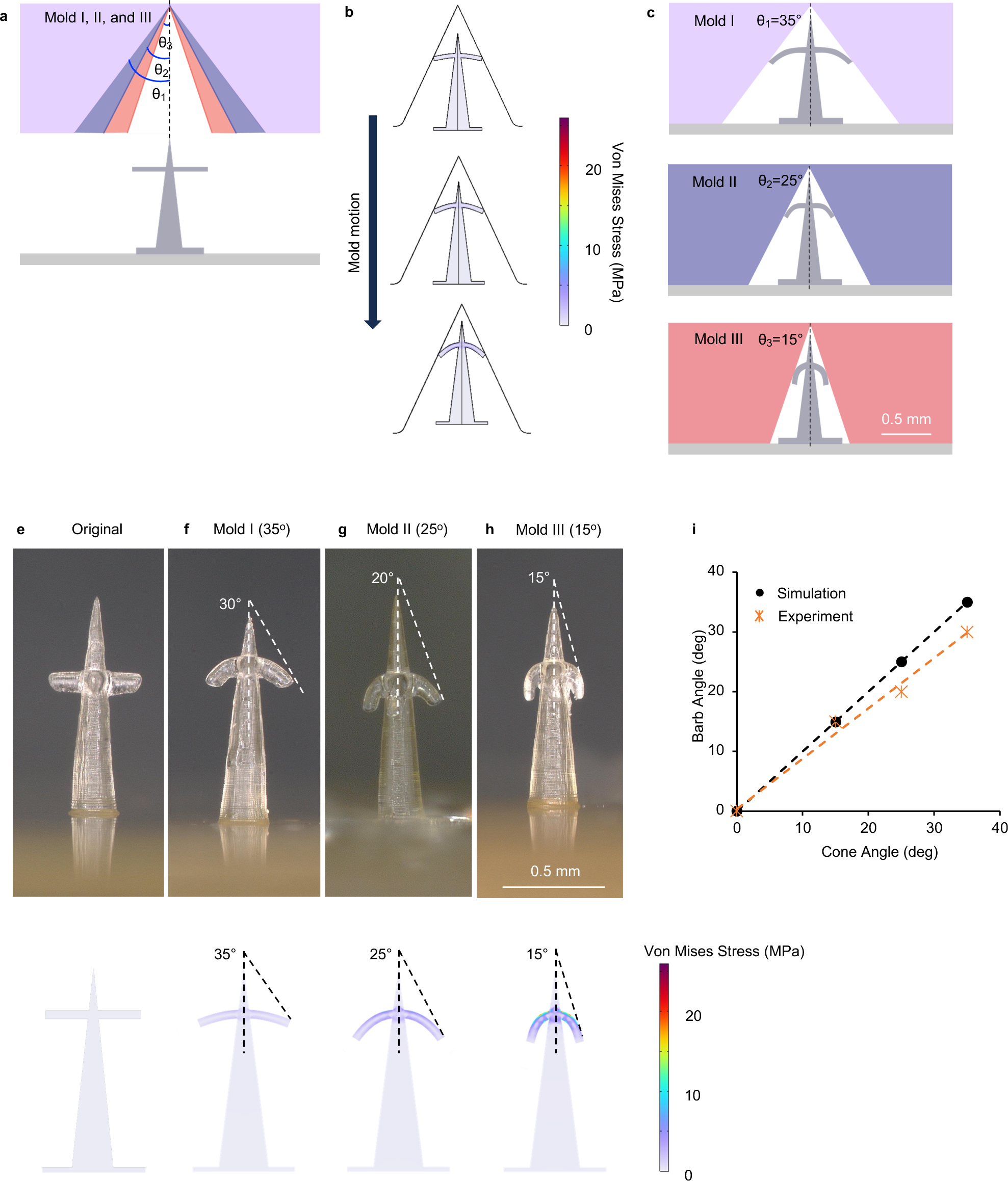
Controlled deformation of backward-facing barbs. **a,** Design of the cone-shaped PDMS mold with different angles. **b,** Numerical simulation of the barb deformation process. **c,** Schematic illustration of barbs deformed by molds of different angles. **e-h,** Experimental and numerical results of barb deformation using molds of different angles (**e,** Original. **f,** Mold I (35°). **g,** Mold II (25°). **h,** Mold III (15°)). **i**, Relationship of the mold angle and barb angle in experiment and simulation.

**Supplementary Fig. 8.**
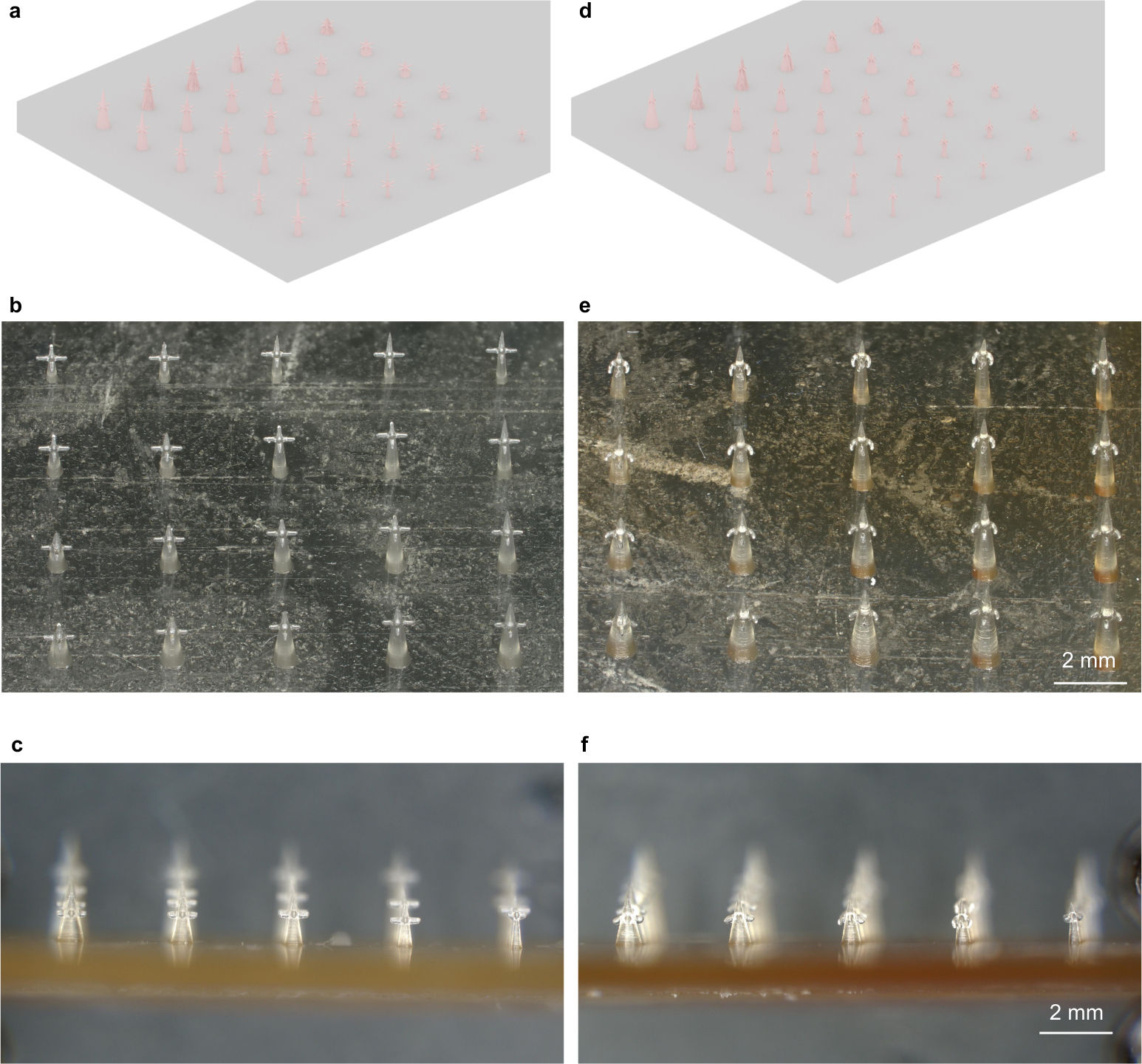
Massive manufacturing of barbed microneedle arrays with microneedles of different dimensions using the deformation method. **a,** 3D rendering of microneedles with horizontal barbs. **b,** Top view of microneedles with horizontal barbs. **c,** Side-view of microneedles with horizontal barbs. **d,** 3D rendering of microneedles after barb deformation. **e,** Top view of microneedles with backward-facing barbs. **f,** Side-view of microneedles with backward-facing barbs.

**Supplementary Fig. 9.**
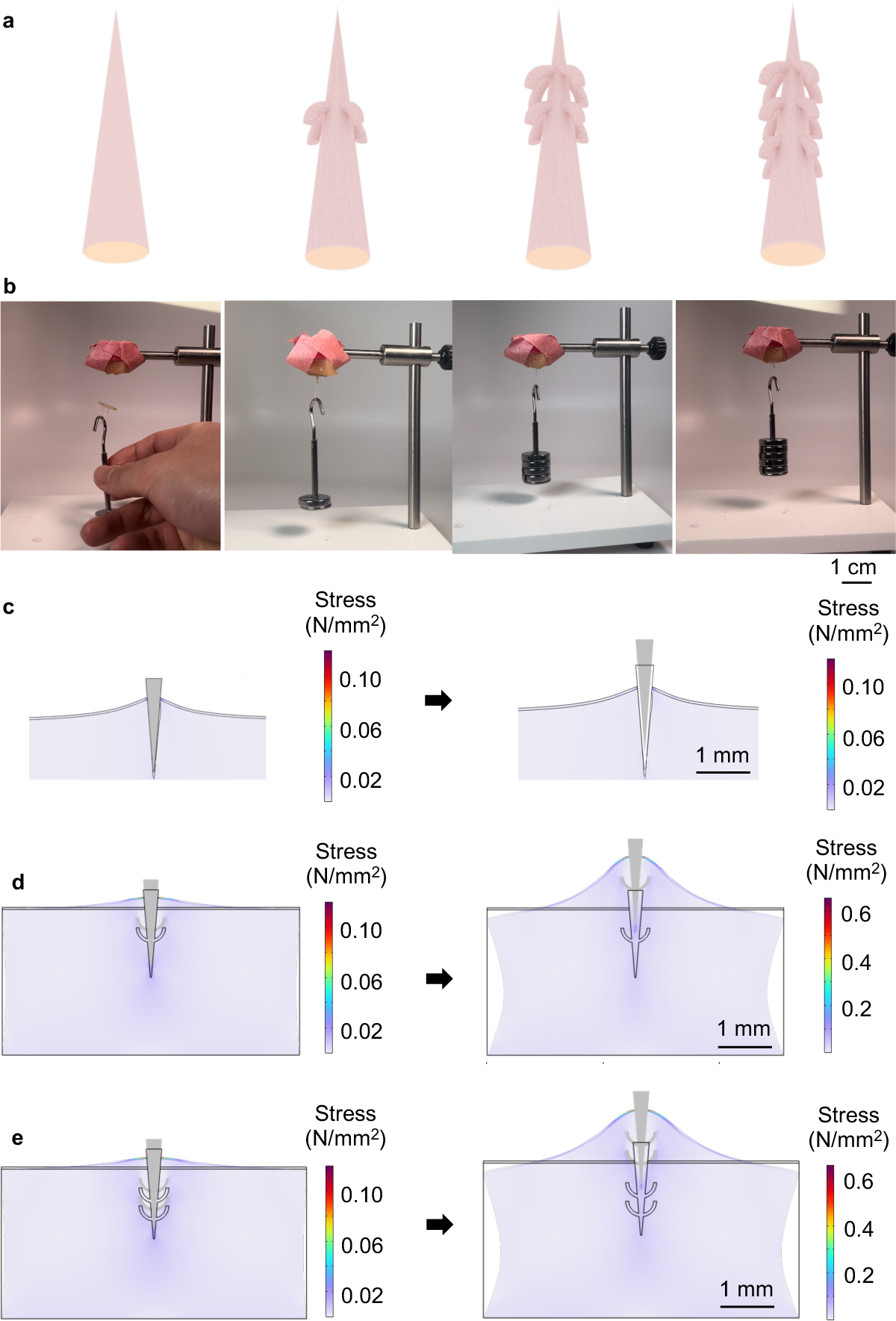
Characterization of tissue adhesion enhanced by backward-facing barbs. **a,** 3D rendering of microneedles with different numbers of rows of barbs. **b,** Experimental results showing the maximum weight withstood by microneedles with 0, 1, 2, and 3 rows of barbs. **c,** Numerical simulation of tissue adhesion of a bare microneedle. **d,** Numerical simulation of tissue adhesion of a microneedle with 1 row of barbs. **e,** Numerical simulation of tissue adhesion of a microneedle with 2 rows of barbs.

**Supplementary Fig. 10.**
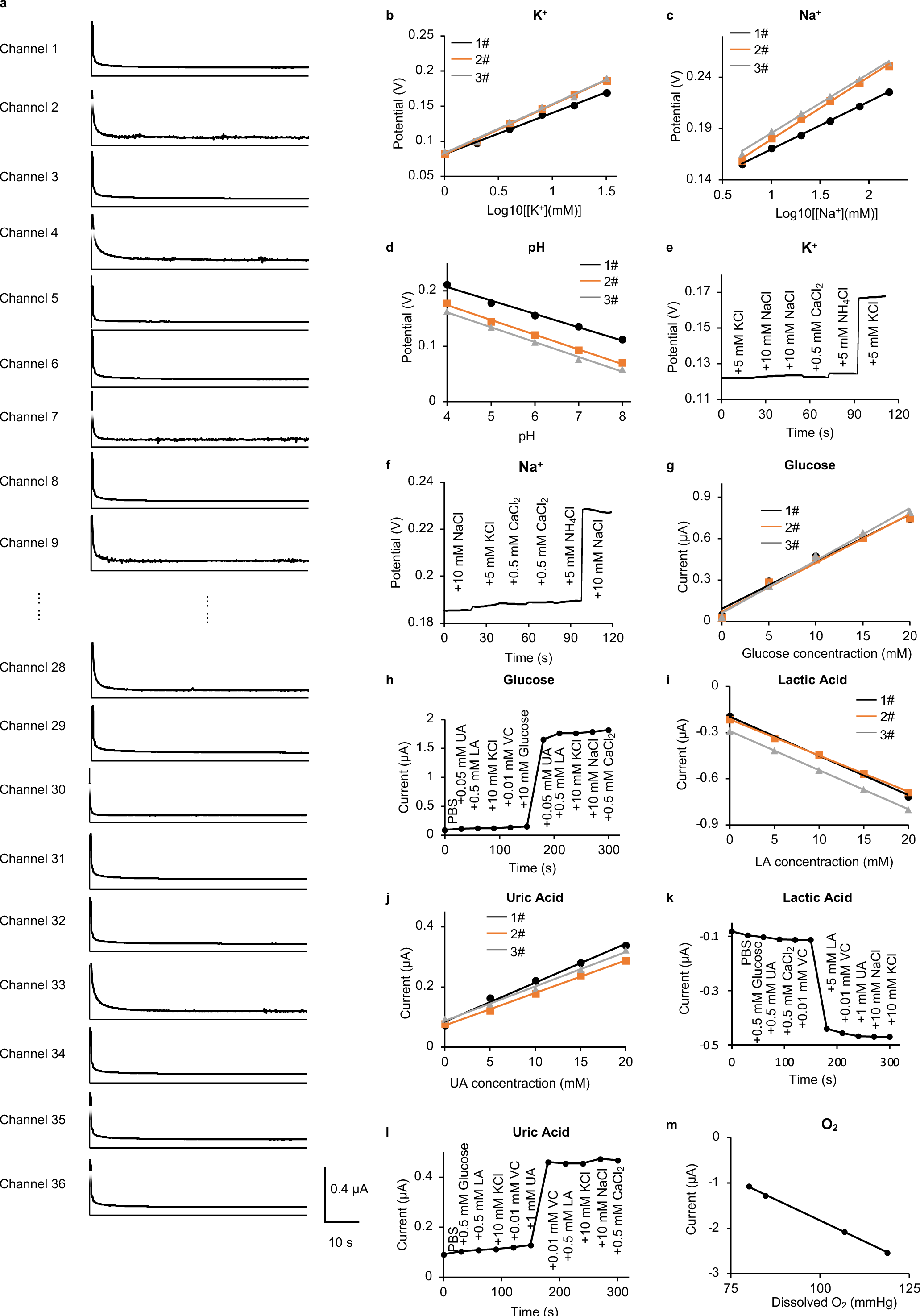
Characterization of electrochemical sensors. **a,** Amperometric response of microneedle glucose sensors in a 6×6 SMART. **b,** Linearity and reproducibility of the K^+^ sensor. **c,** Linearity and reproducibility of the Na^+^ sensor. **d,** Linearity and reproducibility of the pH sensor. **e,** Interference test of the K^+^ sensor. **f,** Interference test of the Na^+^ sensor. **g,** Linearity and reproducibility of the glucose sensor. **h,** Interference test of the glucose sensor. **i,** Linearity and reproducibility of the LA sensor. **j,** Linearity and reproducibility of the UA sensor. **k,** Interference test of the LA sensor. **l,** Interference test of the UA sensor. **m,** Linearity of the oxygen sensor.

**Supplementary Fig. 11.**
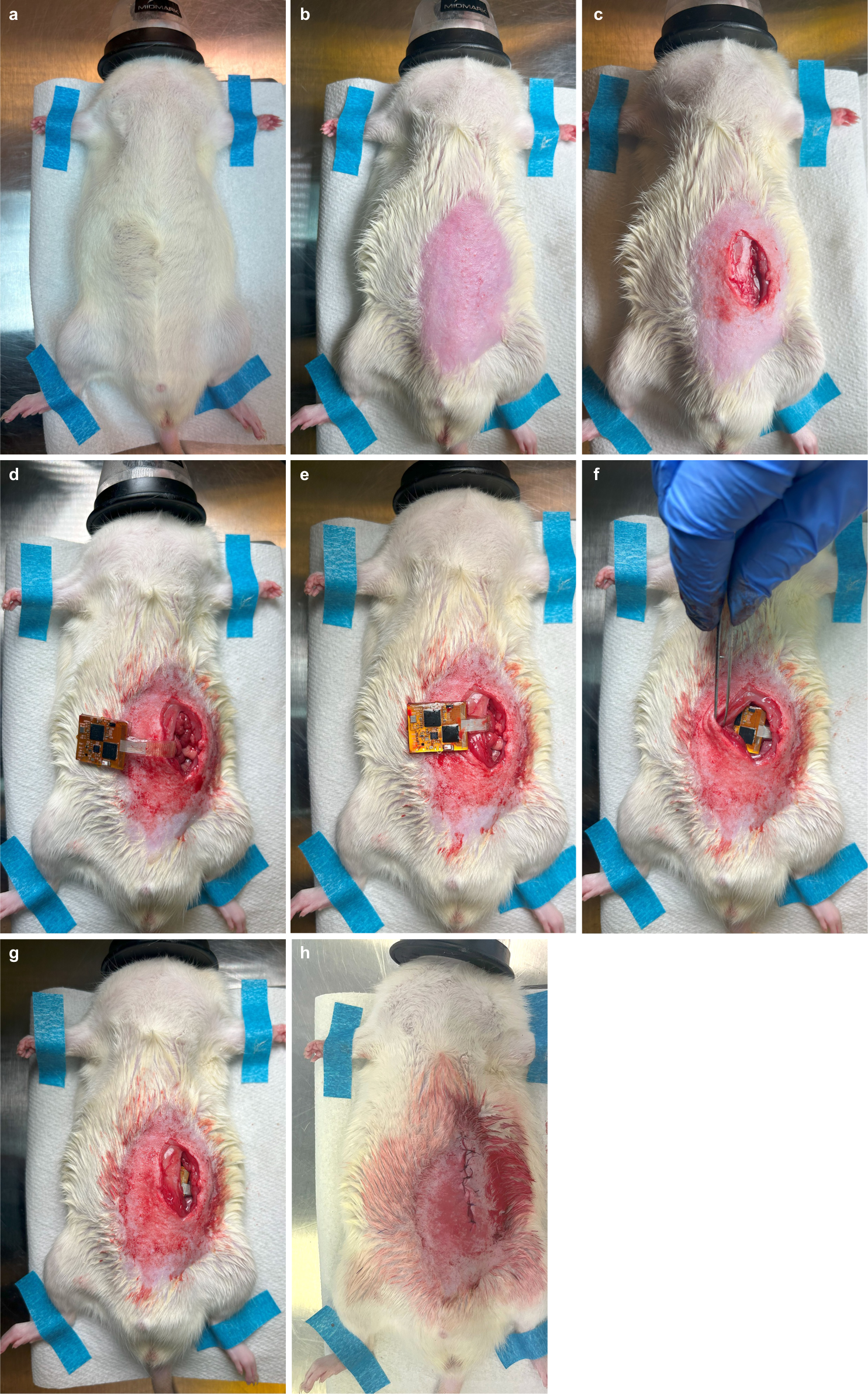
Implantation procedures. **a,** Anesthesia. **b,** Hair removal. **c,** Incision opening. **d,** Mounting the SMART on the target organ. **e,** Insertion of the serpentines. **f,** Insertion of the electronics. **g,** Affixing the electronics to the abdominal wall by tissue adhesive. **h,** Closing the incision.

**Supplementary Fig. 12.**
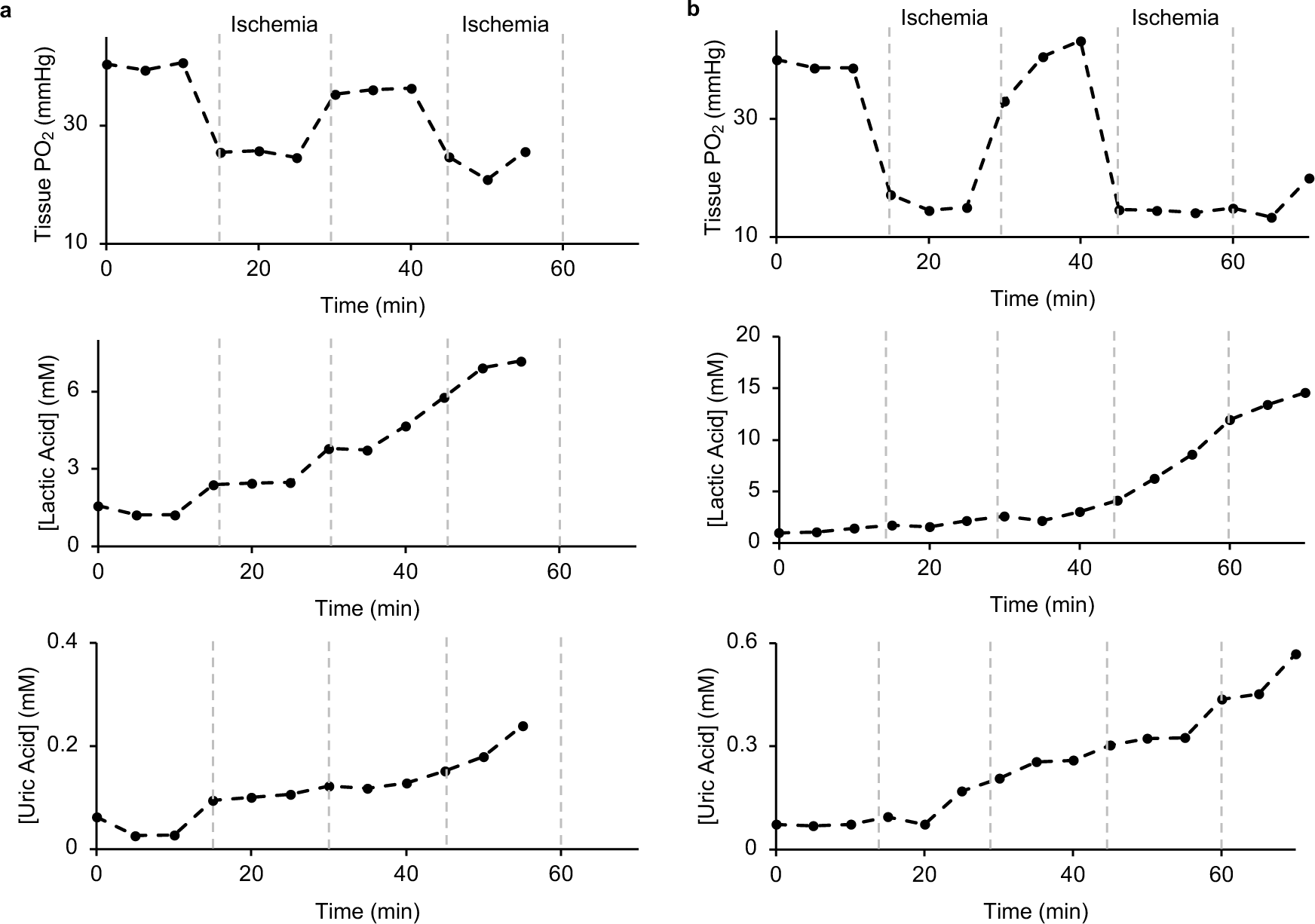
Kidney monitoring results in 2 additional animals. **a,** Additional animal 1. **b,** Additional animal 2.

**Supplementary Fig. 13.**
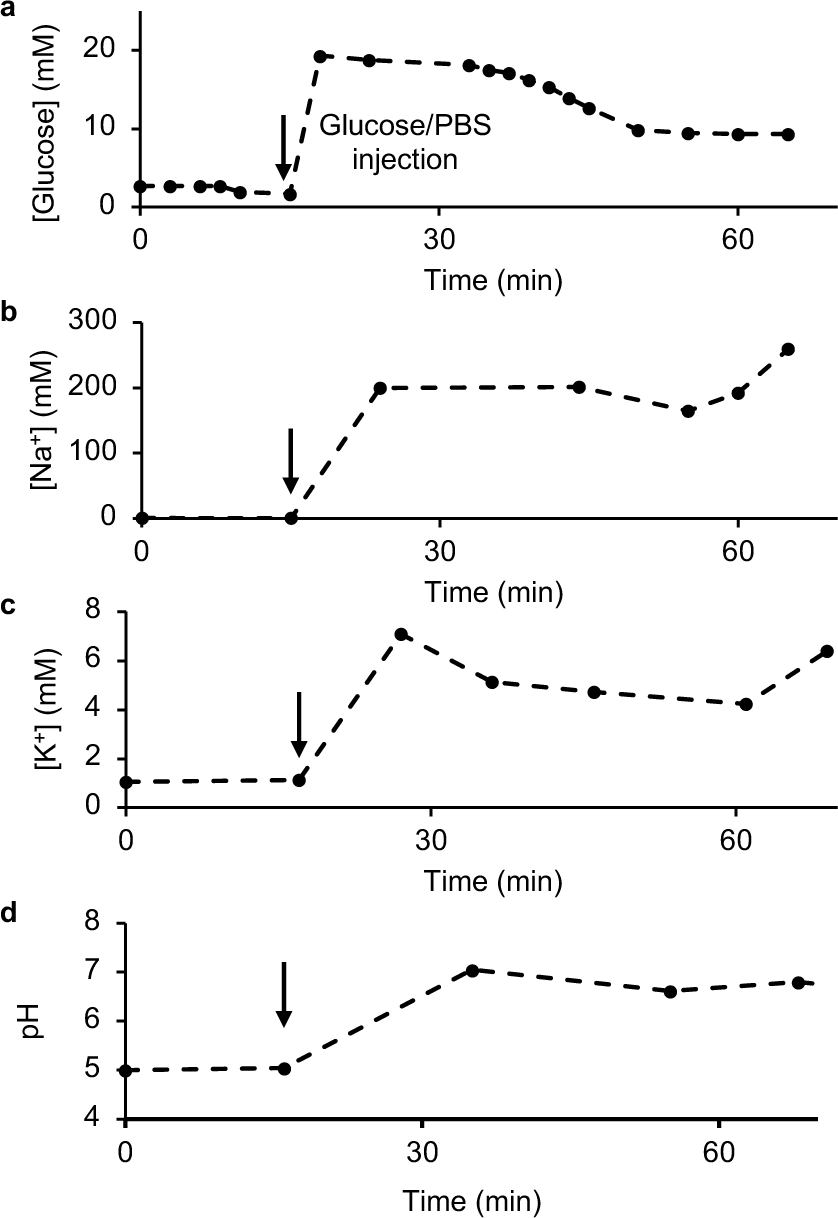
Gut monitoring results in 1 additional animal.

## References

1. Domenghino, A. et al. Consensus recommendations on how to assess the quality of surgical interventions. Nature Medicine 29, 811–822 (2023).

2. Javed, H. et al. Challenges and Solutions in Postoperative Complications: A Narrative Review in General Surgery. Cureus 15, e50942 (2023).

3. Dobson, G.P. Trauma of major surgery: A global problem that is not going away. International Journal of Surgery 81, 47–54 (2020).

4. Ackland, G.L. & Abbott, T.E.F. Hypotension as a marker or mediator of perioperative organ injury: a narrative review. British Journal of Anaesthesia 128, 915–930 (2022).

5. Conrad, C. & Eltzschig, H.K. Disease Mechanisms of Perioperative Organ Injury. 131, 1730–1750 (2020).

6. Gold, A.K., Mandelbaum, T. & Fleisher, L.A. in The High-risk Surgical Patient. (eds. P. Aseni, A.M. Grande, A. Leppäniemi & O. Chiara) 323–343 (Springer International Publishing, Cham; 2023).

7. Bindu, S., Mazumder, S. & Bandyopadhyay, U. Non-steroidal anti-inflammatory drugs (NSAIDs) and organ damage: A current perspective. Biochemical Pharmacology 180, 114147 (2020).

8. Connelly, K. & Morand, E.F. Systemic lupus erythematosus: a clinical update. Internal Medicine Journal 51, 1219–1228 (2021).

9. Ouyang, W. et al. A wireless and battery-less implant for multimodal closed-loop neuromodulation in small animals. Nature Biomedical Engineering 7, 1252–1269 (2023).

10. Prowle, J.R. et al. Postoperative acute kidney injury in adult non-cardiac surgery: joint consensus report of the Acute Disease Quality Initiative and PeriOperative Quality Initiative. Nature Reviews Nephrology 17, 605–618 (2021).

11. Devarbhavi, H. et al. Drug-induced liver injury: Asia Pacific Association of Study of Liver consensus guidelines. Hepatology International 15, 258–282 (2021).

12. Gharbieh, S., Reeves, F. & Challacombe, B. The prostatic middle lobe: clinical significance, presentation and management. Nature Reviews Urology 20, 645–653 (2023).

13. Sah, B.K. et al. Predictive factors and diagnostic significance of CT findings for anastomotic leak after gastric cancer surgery: A retrospective analysis. Aging and Cancer 4, 85–93 (2023).

14. Chung, R. et al. Survival outcomes in patients with muscle invasive bladder cancer undergoing radical vs. partial cystectomy. Urologic Oncology: Seminars and Original Investigations 41, 356.e311–356.e318 (2023).

15. Terrault, N.A., Francoz, C., Berenguer, M., Charlton, M. & Heimbach, J. Liver Transplantation 2023: Status Report, Current and Future Challenges. Clinical Gastroenterology and Hepatology 21, 2150–2166 (2023).

16. Mesnard, B. et al. Kidney transplantation from elderly donors (> 70 years): a systematic review. World Journal of Urology 41, 695–707 (2023).

17. Attieh, R.M. et al. Improved outcomes of kidney after liver transplantation after the implementation of the safety net policy. Liver Transplantation (9900).

18. Guo, H. et al. Wireless implantable optical probe for continuous monitoring of oxygen saturation in flaps and organ grafts. Nature Communications 13, 3009 (2022).

19. Kivimäki, M., Bartolomucci, A. & Kawachi, I. The multiple roles of life stress in metabolic disorders. Nature Reviews Endocrinology 19, 10–27 (2023).

20. Elsenosy, A.M., Hassan, E., Abdelgader, M., Elgamily, O.S. & Hegazy, A. Enhanced Recovery After Surgery (ERAS) Approach: A Medical Complex Experience. Cureus 15, e51208 (2023).

21. Himawan, A. et al. Where Microneedle Meets Biomarkers: Futuristic Application for Diagnosing and Monitoring Localized External Organ Diseases. Advanced Healthcare Materials 12, 2202066 (2023).

22. Lawrence, R., Watters, M., Davies, C.R., Pantel, K. & Lu, Y.-J. Circulating tumour cells for early detection of clinically relevant cancer. Nature Reviews Clinical Oncology 20, 487–500 (2023).

23. Pieper, C.C. Back to the Future II—A Comprehensive Update on the Rapidly Evolving Field of Lymphatic Imaging and Interventions. Investigative Radiology 58 (2023).

24. Morriss, R. et al. Connectivity-guided intermittent theta burst versus repetitive transcranial magnetic stimulation for treatment-resistant depression: a randomized controlled trial. Nature Medicine 30, 403–413 (2024).

25. Reeves, P.T., James-Davis, L.T. & Khan, M.A. Gastrointestinal Bleeding in the Neonate: Updates on Diagnostics, Therapeutics, and Management. NeoReviews 24, e403–e413 (2023).

26. Gedela, M. et al. Mitral Valve Intervention in Elderly or High-Risk Patients: A Review of Current Surgical and Interventional Management. Canadian Journal of Cardiology 40, 250–262 (2024).

27. Ouyang, W. et al. An implantable device for wireless monitoring of diverse physio-behavioral characteristics in freely behaving small animals and interacting groups. Neuron (2024).

28. Jung, Y.H. et al. Injectable Biomedical Devices for Sensing and Stimulating Internal Body Organs. Advanced Materials 32, 1907478 (2020).

29. Zheng, M., Sheng, T., Yu, J., Gu, Z. & Xu, C. Microneedle biomedical devices. Nature Reviews Bioengineering 2, 324–342 (2024).

30. Vora, L.K., et al. Microneedle-based biosensing. Nature Reviews Bioengineering 2, 64–81 (2024).

31. Zhu, D.D. et al. Microneedle-Coupled Epidermal Sensors for In-Situ-Multiplexed Ion Detection in Interstitial Fluids. ACS Applied Materials & Interfaces 15, 14146–14154 (2023).

32. Huang, X. et al. 3D-assembled microneedle ion sensor-based wearable system for the transdermal monitoring of physiological ion fluctuations. Microsystems & Nanoengineering 9, 25 (2023).

33. Li, J., Wei, M. & Gao, B. A Review of Recent Advances in Microneedle-Based Sensing within the Dermal ISF That Could Transform Medical Testing. ACS Sensors 9, 1149–1161 (2024).

34. Lin, S. et al. Wearable microneedle-based electrochemical aptamer biosensing for precision dosing of drugs with narrow therapeutic windows. Science advances 8, eabq4539 (2022).

35. Li, X. et al. A Fully Integrated Closed-Loop System Based on Mesoporous Microneedles-Iontophoresis for Diabetes Treatment. Advanced Science 8, 2100827 (2021).

36. Tehrani, F. et al. An integrated wearable microneedle array for the continuous monitoring of multiple biomarkers in interstitial fluid. Nature Biomedical Engineering 6, 1214–1224 (2022).

37. Huang, S. et al. Petromyzontidae-Biomimetic Multimodal Microneedles-Integrated Bioelectronic Catheters for Theranostic Endoscopic Surgery. Advanced Functional Materials 33, 2214485 (2023).

38. Luo, X., Yang, L. & Cui, Y. Microneedles: materials, fabrication, and biomedical applications. Biomedical Microdevices 25, 20 (2023).

39. Li, S. et al. Microneedle array facilitates hepatic sinusoid construction in a large-scale liver-acinus-chip microsystem. Microsystems & Nanoengineering 9, 75 (2023).

40. Li, J. et al. High-performance flexible microneedle array as a low-impedance surface biopotential dry electrode for wearable electrophysiological recording and polysomnography. Nano-Micro Letters 14, 132 (2022).

41. Wang, Z. et al. Microneedle patch for the ultrasensitive quantification of protein biomarkers in interstitial fluid. Nature Biomedical Engineering 5, 64–76 (2021).

42. Umale, S. et al. Experimental mechanical characterization of abdominal organs: liver, kidney & spleen. Journal of the Mechanical Behavior of Biomedical Materials 17, 22–33 (2013).

43. Tran, P., Lau, C., Joshi, M., Maddock, H. & Banerjee, P. BS11 Changes in myocyte architecture, contractile function and energetics in patients with heart failure before and after ventricular assist device implantation as a sign of myocardial recovery: a systematic review and meta-analysis. Heart 109, A252–A254 (2023).

44. Steenackers, N. et al. Understanding the gastrointestinal tract in obesity: From gut motility patterns to enzyme secretion. Neurogastroenterology & Motility 36, e14758 (2024).

45. Jang, T.-M. et al. Expandable and implantable bioelectronic complex for analyzing and regulating real-time activity of the urinary bladder. Science advances 6, eabc9675 (2020).

46. Ji, H. et al. Skin-integrated, biocompatible, and stretchable silicon microneedle electrode for long-term EMG monitoring in motion scenario. npj Flexible Electronics 7, 46 (2023).

47. Kim, H. et al. Skin preparation–free, stretchable microneedle adhesive patches for reliable electrophysiological sensing and exoskeleton robot control. Science Advances 10, eadk5260 (2024).

48. Zhao, Q. et al. Highly stretchable and customizable microneedle electrode arrays for intramuscular electromyography. Science Advances 10, eadn7202 (2024).

49. Yang, S.Y. et al. A bio-inspired swellable microneedle adhesive for mechanical interlocking with tissue. Nature Communications 4, 1702 (2013).

50. Han, D. et al. 4D printing of a bioinspired microneedle array with backward-facing barbs for enhanced tissue adhesion. Advanced Functional Materials 30, 1909197 (2020).

51. Farzam, M., Beitollahpoor, M. & Pesika, N.S. Nature-Inspired Directional Microneedle Structures for Reversible Gripping on Skin and Fibrous Materials. Advanced Engineering Materials 26, 2400149 (2024).

52. Chen, Z. et al. Additive manufacturing of honeybee-inspired microneedle for easy skin insertion and difficult removal. ACS applied materials & interfaces 10, 29338–29346 (2018).

53. Wang, Y. et al. Digital automation of transdermal drug delivery with high spatiotemporal resolution. Nature Communications 15, 511 (2024).

54. Malek-Khatabi, A. et al. Recent progress in PLGA-based microneedle-mediated transdermal drug and vaccine delivery. Biomaterials Science 11, 5390–5409 (2023).

55. Shen, J. & Burgess, D.J. Accelerated in vitro release testing of implantable PLGA microsphere/PVA hydrogel composite coatings. International Journal of Pharmaceutics 422, 341–348 (2012).

56. Song, Y. et al. 3D-printed epifluidic electronic skin for machine learning–powered multimodal health surveillance. Science Advances 9, eadi6492 (2023).

57. Wang, J. Electrochemical Glucose Biosensors. Chemical Reviews 108, 814–825 (2008).

58. Rivas, L. et al. Micro-needle implantable electrochemical oxygen sensor: ex-vivo and in-vivo studies. Biosensors and Bioelectronics 153, 112028 (2020).

59. Gerwig, R. et al. PEDOT–CNT Composite Microelectrodes for Recording and Electrostimulation Applications: Fabrication, Morphology, and Electrical Properties. Frontiers in Neuroengineering 5 (2012).

60. Buckthorpe, M.W., Hannah, R., Pain, T.G. & Folland, J.P. Reliability of neuromuscular measurements during explosive isometric contractions, with special reference to electromyography normalization techniques. Muscle & Nerve 46, 566–576 (2012).

61. Yang, B., Fung, A., Pac-Soo, C. & Ma, D. Vascular surgery-related organ injury and protective strategies: update and future prospects. BJA: British Journal of Anaesthesia 117, ii32–ii43 (2016).

62. Madhvapathy, S.R. et al. Implantable bioelectronic systems for early detection of kidney transplant rejection. Science 381, 1105–1112 (2023).

63. Tasoulis, M.K. & Douzinas, E.E. Hypoxemic reperfusion of ischemic states: an alternative approach for the attenuation of oxidative stress mediated reperfusion injury. Journal of Biomedical Science 23, 1–8 (2016).

64. Guyton, K. & Alverdy, J.C. The gut microbiota and gastrointestinal surgery. Nature Reviews Gastroenterology & Hepatology 14, 43–54 (2017).

65. Li, S. et al. Bioresorbable, wireless, passive sensors for continuous pH measurements and early detection of gastric leakage. Science Advances 10, eadj0268 (2024).

66. Lu, D. et al. Bioresorbable wireless sensors as temporary implants for in vivo measurements of pressure. Advanced Functional Materials 30, 2003754 (2020).

67. Choi, H.J. et al. MG-63 osteoblast-like cell proliferation on auxetic PLGA scaffold with mechanical stimulation for bone tissue regeneration. Biomaterials research 20, 33 (2016).

